# The role of REM sleep in neural differentiation of memories in the hippocampus

**DOI:** 10.1101/2024.11.01.621588

**Authors:** Elizabeth A. McDevitt, Ghootae Kim, Nicholas B. Turk-Browne, Kenneth A. Norman

## Abstract

When faced with a familiar situation, we can use memory to make predictions about what will happen next. If such predictions turn out to be erroneous, the brain can adapt by differentiating the representations of the cues that generated the prediction from the mispredicted item itself, reducing the likelihood of future prediction errors. Prior work by Kim et al. (2017) found that violating a sequential association in a statistical learning paradigm triggered differentiation of the neural representations of the associated items in the hippocampus. Here, we used fMRI to test the preregistered hypothesis that this hippocampal differentiation occurs only when violations are followed by rapid eye movement (REM) sleep. In the morning, participants first learned that some items predict others (e.g., A predicts B) then encountered a violation in which a predicted item (B) failed to appear when expected after its associated item (A); the predicted item later appeared on its own after an unrelated item. Participants were then randomly assigned to one of three conditions: remain awake, take a nap containing non-REM sleep only, or take a nap with both non-REM and REM sleep. While the predicted results were not observed in the preregistered left CA2/3/DG ROI, we did observe evidence for our hypothesis in closely related hippocampal ROIs, uncorrected for multiple comparisons: In right CA2/3/DG, differentiation in the group with REM sleep was greater than in the groups without REM sleep (wake and non-REM nap); this differentiation was item-specific and concentrated in right DG. Differentiation effects were also greater in bilateral DG when the predicted item was more strongly reactivated during the violation. Overall, the results presented here provide initial evidence linking REM sleep to changes in the hippocampal representations of memories in humans.

## Introduction

When we retrieve a memory, related memories often come to mind. In some situations, this may be helpful: for example, you enter a familiar environment and can predict who or what you will encounter. But what if your prediction is wrong, and instead becomes a source of interference for memory retrieval? One way the brain might mitigate prediction errors is by adaptively disconnecting the mispredicted item from its old context and binding it to a new, updated context, effectively pushing a mispredicted memory away from its old cue in representational space (i.e., neural differentiation). Here, we investigate whether this process of prediction-based neural differentiation is supported by a period of optimized memory consolidation that includes sleep.

Our work builds on a prior functional MRI (fMRI) study by Kim et al. (2017) that found that prediction errors lead to neural differentiation in the hippocampus. Specifically, they found that — when an item predicted in a particular context (e.g., A predicts B) failed to appear and was later restudied in a different context — the neural representations of A and B became less similar in the left CA2/3/DG subregion of the hippocampus. Kim et al. (2017) explained this result in terms of an unuspervised learning principle called the nonmonotonic plasticity hypothesis (NMPH) (Ritvo et al., 2019), which posits a U-shaped relationship between the coactivation of two memories (A and B) and learning; according to the NMPH, strong coactivation of the B memory while retrieving the A memory will lead to strengthening of the connections between A and B, moderate coactivation of the B memory will lead to synaptic weakening, and little or no activation of B will lead to no change in the synaptic connections between A and B (Detre et al., 2013; Newman and Norman, 2010). Kim et al. (2017) argued that, when A predicts B but B does not appear, this unconfirmed prediction leads to moderate activation of B, which — according to the NMPH — weakens the connections between the unique features of B and the features it formerly shared with A (for further evidence that B is weakened, see Kim et al., 2014); when item B is presented on a subsequent trial, it then activates a different set of features (not shared with A) and incorporates these new features into its neural representation (Hulbert and Norman, 2015; Ritvo et al., 2019, 2024). The overall effect of this process is neural differentiation — decreased overlap in the populations of neurons that encode A and B.

An important detail of the Kim et al. (2017) study (and some other fMRI studies that have found differentiation (e.g., Favila et al., 2016) is that the interval between learning and the final measurement of neural representations contained a night of sleep, raising the question of how offline consolidation processes might contribute to the observed representational changes. Numerous studies have found that neural activity is reactivated (i.e., replayed) during sleep (Wilson and McNaughton, 1994; Louie and Wilson, 2001; Ji and Wilson, 2007), and this is thought to be a critical mechanism underlying sleep-dependent memory consolidation (Diekelmann and Born, 2010; Klinzing et al., 2019). Neural network modeling suggests that rapid eye movement (REM) sleep, in particular, serves as a focused period of interleaved replay of related memories (Norman et al., 2005; Singh et al., 2022; Guerreiro and Clopath, 2024). During REM sleep, brain activity is not guided by environmental stimuli, and the hippocampus and cortex are relatively uncoupled (Cantero et al., 2003; Diekelmann and Born, 2010). This allows the hippocampus (and cortex) to autonomously rehearse stored memories in an unsupervised, interleaved manner, meaning that spreading activation within each network selects the ‘targets’ for learning, while coactivating related memories in the process. This is in contrast to non-REM (NREM) sleep, when neural activity between the hippocampus and cortex is tightly coupled and the hippocampus is replaying information to cortex (Diekelmann and Born, 2010). One way to think about this is that, during NREM sleep, the hippocampus is busy training cortex; during REM sleep, the hippocampus is freed of that obligation and can focus on fine-tuning its own network of information. This led us to hypothesize that the brain identifies memories vulnerable to prediction errors during wake, and then implements the restructuring needed to address those prediction errors during sleep. Specifically, we hypothesized that REM sleep should be the critical sleep stage driving hippocampal neural differentiation.

In a preregistered study, we tested this hypothesis in a day-long experiment using fMRI to measure how neural representations differentiate across periods of wake and sleep. We used the same task and the same general pre/post design as Kim et al. (2017). In the morning, we first obtained pre-learning fMRI “snapshots” of each item’s initial neural representation by showing participants all relevant items and extracting the spatial pattern of BOLD activity corresponding to each item. Participants then completed the learning task that was previously shown to induce neural differentiation (Kim et al., 2017). Next, participants were randomly assigned to one of three EEG-recorded offline conditions: a nap composed of NREM sleep only (NREM), a nap with both NREM and REM sleep (REM), or a period of quiet wakefulness (Wake). Later the same day, we obtained post-learning (and post-sleep, in the case of the nap groups) fMRI snapshots of each item’s neural representation. Comparing pre- and post-learning snapshots allows us to assess how the intervening task and subsequent sleep or wake conditions altered the neural representations.

In our preregistration, the overarching prediction was that violations during learning (i.e., instances when A predicted B but B did not appear), followed by REM sleep—versus not having REM sleep (i.e., Wake or NREM only)—would lead to neural differentiation of A and B. Kim et al. (2017) also found that — when A predicted B, but B did not appear — stronger activation of memory B predicted higher levels of (subsequent) neural differentiation in the hippocampus; they explained this in terms of low activation being associated with no synaptic change and higher (moderate) levels of activation being associated with synaptic weakening and, through this, differentiation. We also tested for this relationship in our study, predicting that it would be more evident in the REM group than in the other two groups. Because the Kim et al. (2017) study found these differentiation effects in the left CA2/3/dentate gyrus (DG) subfield of the hippocampus, we focused initially on this specific subregion.

As described below, we did not find evidence for the predicted effects in the left CA2/3/DG, but we did find the predicted pattern of results (uncorrected for multiple comparisons) in the same hippocampal subfield, lateralized to the right hemisphere. Compared to the Wake and NREM groups, the REM group showed more neural differentiation in the right CA2/3/DG (and right DG alone). Additionally, there was a stronger relationship between B activation and differentiation in bilateral DG in the REM group. Together, these findings provide support for our hypothesis that REM sleep facilitates learning-dependent representational change in the hippocampus, although more work is needed to firmly establish this point and to understand whether hemispheric differences are real or reflect noisy fMRI measurements from small and highly precise anatomical regions.

## Results

An overview of the study is illustrated in Figure 1. In session 1, participants viewed streams of scene and face images with embedded regularities while performing a cover task in the MRI scanner. The streams were composed of 96 individual scene pairs (split into six runs), where each pair had scene A as the first item and a different scene B as the second item; as is typical of statistical learning studies, participants were not made aware of this pair structure. Each A-B pair (A followed by B) was shown three times, inserted in the stream continuously amongst the other pairs, creating the expectation that B would follow A. After three repetitions of a given pair, critical trials created two within-subject task conditions: violation and nonviolation (48 pairs each). For violation pairs, there were two “violation” events in which B failed to follow A. Instead, the sequence was A-X and A-Y, where X and Y were novel faces. Following each violation event, the B item was subsequently “restudied” on its own, meaning it appeared in a novel context, not preceded by its pairmate A. Pairs in the nonviolation condition also had three initial repetitions, but no violation events. To match the frequency of B item exposures in the two conditions, the nonviolation condition also included two B “restudy” events. At the beginning of each run, the A and B members of each pair were shown once separately (i.e., B did not follow A) and randomly intermixed with scenes from other pairs. This first presentation of each scene was used to estimate each scene’s neural representation (“pre-learning snapshot”), uncontaminated by its pairmate. Following session 1, participants either remained awake, took a nap with NREM sleep only, or took a nap with NREM and REM sleep. In session 2, participants re-entered the MRI scanner and completed three separate tasks in the following order: 1) one run of “post-learning snapshots” for B scenes; 2) a reward association task (see Supplementary Methods); and 3) one run of post-learning snapshots for X and Y faces.

**Figure 1.**
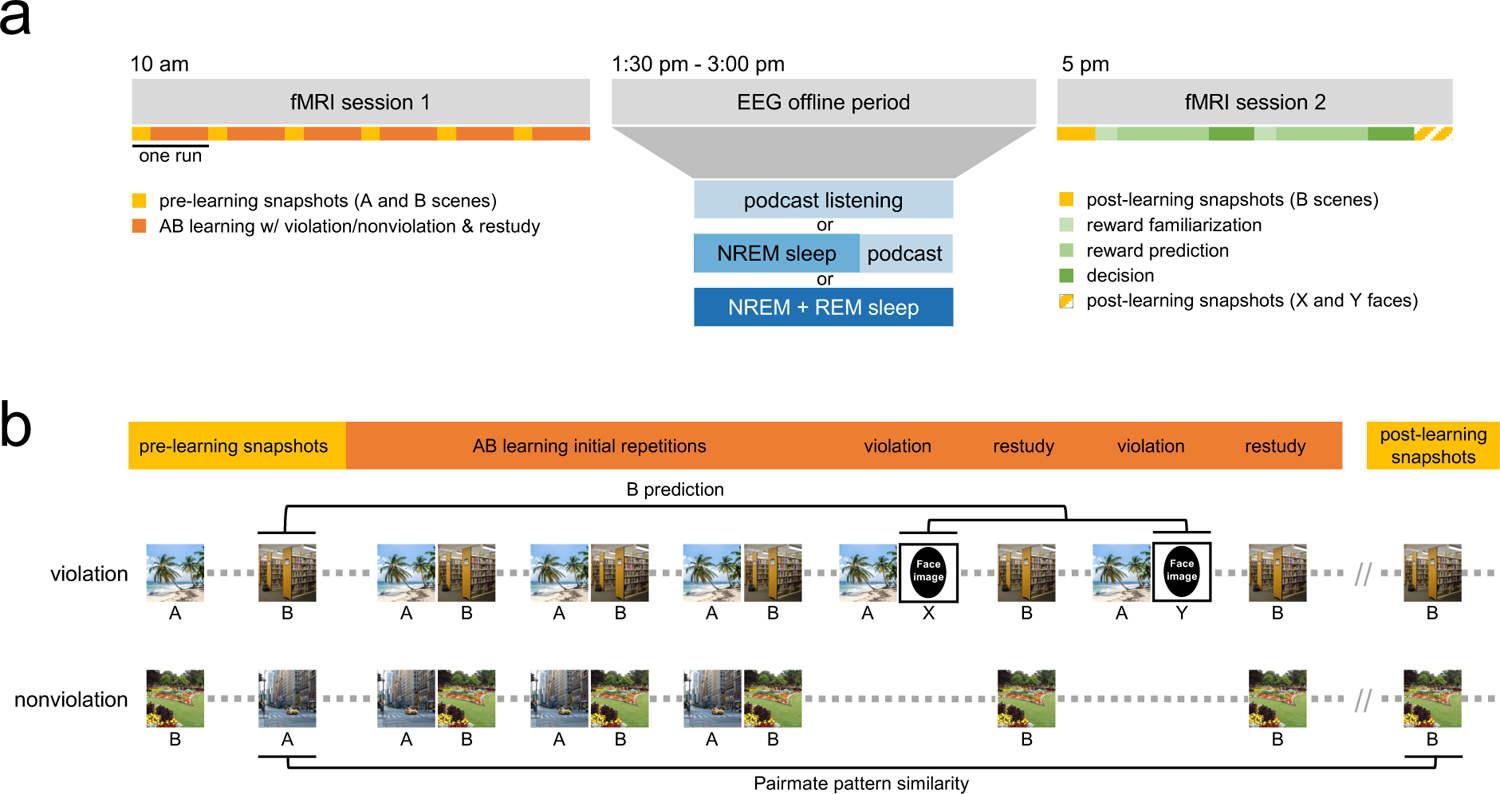
Experimental design and methods. (**a**) Study day timeline. Participants entered the MRI scanner at 10:00 AM and completed an incidental encoding task. Next, participants were randomly assigned to one of three EEG-recorded offline conditions: wake, a short 50-minute nap followed by podcast listening, or a long 90-minute nap. Participants re-entered the MRI scanner at 5:00 PM and completed one run of post-learning “B” scene snapshots, the reward association task, and one run of post-learning “X and Y” face snapshots. **(b)** Task design and analysis schematic. Participants viewed streams of scene and faces images presented one at a time. At the beginning of each run, the A and B members of each pair were shown once separately (i.e., B did not follow A) to obtain pre-learning snapshots of each item. During the learning phase, pairs in the violation condition followed a sequence of three initial learning repetitions followed by two cycles of violation and restudy trials (AB-AB-AB-AX-B-AY-B). The nonviolation control condition did not have any violation events, but did include two B restudy trials (AB-AB-AB-B-B). Each pair’s trials were interleaved with repetitions of other pairs (represented here as gray dots). In session 2, the B scenes were presented one more time in a random order to take post-learning snapshots. To measure neural differentiation, we correlated voxel patterns for the pre-learning snapshot of A and post-learning snapshot of B (“pairmate pattern similarity”) for all pairs. To measure the amount of “B prediction” on violation trials, we correlated the pre-learning snapshot of B and the pattern of activity evoked by the X and Y violation events, then averaged these values for one B prediction score per pair in the violation condition. Note: The scene images in this figure were sourced from the internet and not used as actual stimuli in our experiment; face stimuli have been replaced with black ovals.

### Regions of interest (ROIs)

We were specifically interested in measuring neural differentiation in the hippocampus (Hulbert and Norman, 2015; Schlichting et al., 2015; Chanales et al., 2017; Molitor et al., 2021; Dimsdale-Zucker et al., 2018; Wanjia et al., 2021; Zeithamova et al., 2018; Wammes et al., 2022; Fernandez et al., 2023). Within the hippocampus, we expected to observe differentiation in the left CA2/3/DG subfield, which was the locus of representational change in the Kim et al. (2017) study. We also preregistered right and bilateral CA2/3/DG, and left, right and bilateral CA1 as other regions of interest (ROIs). Neural activity is sparser in CA2/3/DG compared to CA1, making it more difficult for competing memories to come to mind strongly (Schapiro et al., 2017b; Barnes et al., 1990; West et al., 1991; GoodSmith et al., 2017). According to the NMPH, lower levels of memory activation in CA2/3/DG should bias this region toward showing differentiation, whereas higher levels of activation in CA1 should bias this region toward showing integration (Ritvo et al., 2019, 2024). In line with this, most fMRI studies of differentiation that have looked at hippocampal subfields have found that differentiation effects tend to be localized in CA2/3/DG rather than in CA1 (e.g., Dimsdale-Zucker et al., 2018; Wanjia et al., 2021; Molitor et al., 2021; Wammes et al., 2022; but see Zheng et al., 2021). Thus, based on both theoretical grounds (relating to the NMPH) and empirical precedent, there are strong reasons to expect that differentiation effects in our study will be more readily observed in CA2/3/DG than in CA1.

### Behavioral cover task

In session 1, participants completed six runs of incidental encoding while performing a subcategory judgment task (“Is the scene indoor or outdoor?” or “Is the face male or female?”). Each run consisted of pre-learning snapshot trials, three repetitions of A-B pairings, violation and restudy events. In session 2, participants completed two runs of the same subcategory judgment task to obtain post-learning snapshots of scenes and faces.

Session 1 task performance was highly accurate (overall mean = 0.95, sd = 0.04), and did not differ between the three groups (Wake, NREM, REM; F_2,25.02_ = 0.51, p = 0.61). There was no interaction between trial type (pre-learning, A-B pairings, violation events, restudy events) and group (F_6,27.62_ = 0.55, p = 0.77; Figure S1a). Session 2 post-learning snapshot performance was also highly accurate (overall mean = 0.97, sd = 0.02) and did not differ between the three groups (F_2,27.37_ = 0.84, p = 0.44; S1b).

### Nap sleep architecture

Nap sleep architecture variables are summarized in Table 1 and Figure S2. By design, the REM group had significantly greater total sleep time (TST) and minutes of REM sleep than the NREM group (Wilcoxon rank sum test, both *p*s < 0.001). The REM group also had significantly more minutes of Stage 1 sleep (Wilcoxon rank sum test, p = 0.03), Stage 2 sleep (Wilcoxon rank sum test, p < 0.001), and wake after sleep onset (WASO; Wilcoxon rank sum test, p = 0.004). However, the groups did not differ on these values as a percentage of TST (Stage 1 percent: Wilcoxon rank sum test, p = 0.93; Stage 2 percent: Welch’s *t*-test, t_32.5_ = –0.11, p = 0.91; WASO percent: Wilcoxon rank sum test, p = 0.06). There was no difference between groups in minutes of Stage 3 (t_42_ = –0.48, p = 0.64), but the NREM group had a greater percentage of time spent in Stage 3 than the REM group (Welch’s *t*-test, t_28.5_ = 2.55, p = 0.02). The groups did not differ in sleep efficiency (TST/time spent in bed; Wilcoxon rank sum test, p = 0.99).

**Table 1.**
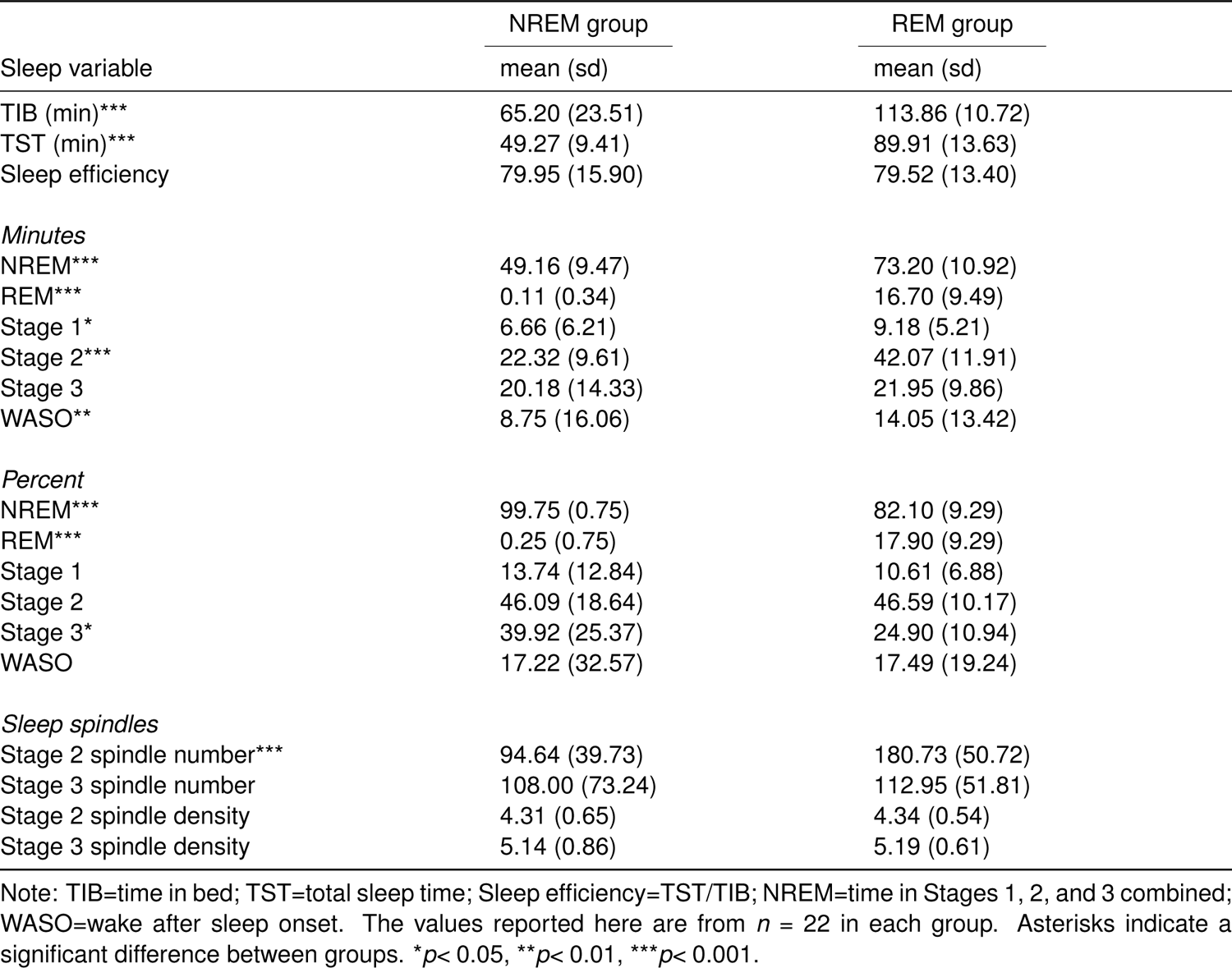
Sleep architecture.

| Sleep variable | NREM group | REM group |
| --- | --- | --- |
|  | mean (sd) | mean (sd) |
| TIB (min)*** | 65.20 (23.51) | 113.86 (10.72) |
| TST (min)*** | 49.27 (9.41) | 89.91 (13.63) |
| Sleep efficiency | 79.95 (15.90) | 79.52 (13.40) |
| *Minutes* |  |  |
| NREM*** | 49.16 (9.47) | 73.20 (10.92) |
| REM*** | 0.11 (0.34) | 16.70 (9.49) |
| Stage 1* | 6.66 (6.21) | 9.18 (5.21) |
| Stage 2*** | 22.32 (9.61) | 42.07 (11.91) |
| Stage 3 | 20.18 (14.33) | 21.95 (9.86) |
| WASO** | 8.75 (16.06) | 14.05 (13.42) |
| *Percent* |  |  |
| NREM*** | 99.75 (0.75) | 82.10 (9.29) |
| REM*** | 0.25 (0.75) | 17.90 (9.29) |
| Stage 1 | 13.74 (12.84) | 10.61 (6.88) |
| Stage 2 | 46.09 (18.64) | 46.59 (10.17) |
| Stage 3* | 39.92 (25.37) | 24.90 (10.94) |
| WASO | 17.22 (32.57) | 17.49 (19.24) |
| *Sleep spindles* |  |  |
| Stage 2 spindle number*** | 94.64 (39.73) | 180.73 (50.72) |
| Stage 3 spindle number | 108.00 (73.24) | 112.95 (51.81) |
| Stage 2 spindle density | 4.31 (0.65) | 4.34 (0.54) |
| Stage 3 spindle density | 5.14 (0.86) | 5.19 (0.61) |

Note: TIB=time in bed; TST=total sleep time; Sleep efficiency=TST/TIB; NREM=time in Stages 1, 2, and 3 combined; WASO=wake after sleep onset. The values reported here are from $n = 22$ in each group. Asterisks indicate a significant difference between groups. \* $p < 0.05$ , \*\* $p < 0.01$ , \*\*\* $p < 0.001$ .

In keeping with the overall greater amount of time spent in Stage 2 sleep, the REM group also had significantly more discrete Stage 2 spindle events (Wilcoxon rank sum test, p < 0.001), but there was no difference in Stage 2 spindle density (spindles/minute) between groups (t_42_ = –0.16, p = 0.87). The groups also did not differ in the number or density of sleep spindles during Stage 3 (Wilcoxon rank sum test, spindle number: p = 0.62; spindle density: p = 0.22).

### Neural differentiation as a function of sleep

To examine how much the neural representation of B items moved away from their A pairmates, we used the same procedure as Kim et al. (2017). We correlated voxel patterns for the pre-learning snapshot of A and post-learning snapshot of B (preA/postB) for all AB pairs within each task condition (violation and nonviolation) for each participant (Figure 1b). Next, we computed the average pattern similarity value for each task condition (Figure S3; note that numerically smaller pattern similarity values indicate less neural overlap, or *more* neural differentiation). Since our general hypothesis was that pattern similarity should be lower for violation compared to nonviolation pairs (Kim et al., 2017), we computed the difference of the violation and nonviolation conditions in each subject. We will refer to this difference as the neural differentiation score; negative values indicate more violation-related neural differentiation.

Our main preregistered hypothesis was that violation-related neural differentiation should be greatest in the REM group, specifically in the left CA2/3/DG subfield of the hippocampus. As the most direct test of this hypothesis, we ran a planned contrast (contrast weights: REM: −1, NREM: 0.5, Wake: 0.5) to determine if having REM sleep, compared to not having REM sleep (i.e., NREM or Wake), resulted in a significantly more negative neural differentiation score. This contrast was not significant in left CA2/3/DG (t_66_ = 0.027, p = 0.98, d = 0.007; Figure 2). We ran the same contrast in our five remaining ROIs (right and bilateral CA2/3/DG and left, right, and bilateral CA1). The predicted pattern of results was present in right CA2/3/DG (t_66_ = 2.19, p = 0.03, d = 0.54), although it was not significant after correcting for multiple comparisons (adjusted alpha = 0.008). The contrast was not significant in bilateral CA2/3/DG (t_66_ = 1.12, p = 0.26, d = 0.28) or any CA1 ROI (left CA1: t_66_ = –0.35, p = 0.73, d = –0.08; right CA1: t_66_ = –0.83, p = 0.41, d = –0.20; bilateral CA1: t_66_ = –0.81, p = 0.42, d = –0.20).

**Figure 2.**
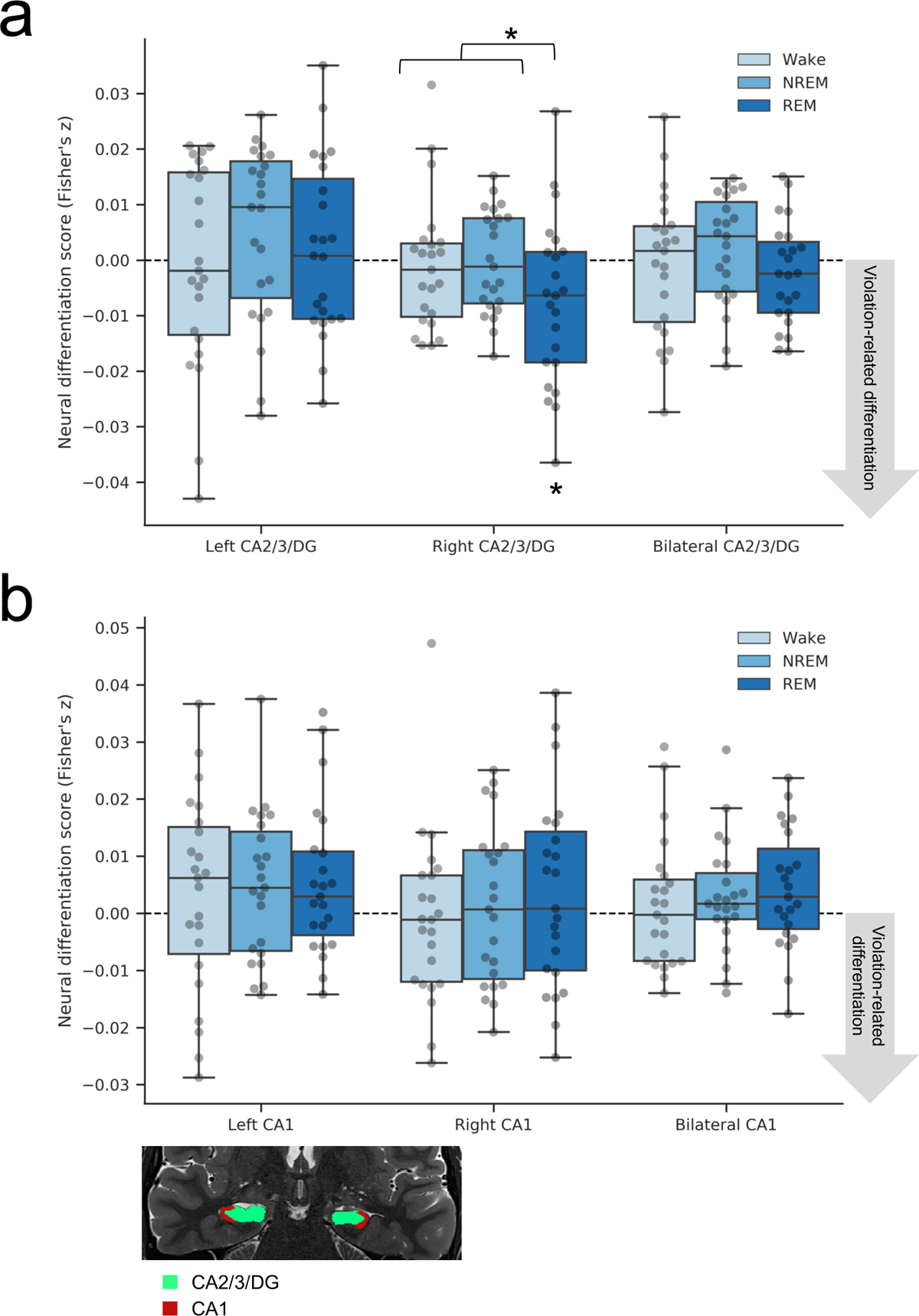
Neural differentiation. Neural differentiation scores were calculated as the difference in preA/postB pattern similarity for the violation minus nonviolation task conditions, in each sleep group in (**a**) CA2/3/DG and (**b**) CA1. A planned contrast in our 6 ROIs revealed more violation-related neural differentiation in the REM group than the Wake and NREM groups in right CA2/3/DG (*p* = 0.03, uncorrected for multiple comparisons). Within the REM group, post-hoc tests showed that the neural differentiation score in right CA2/3/DG was significantly negative (*p* = 0.01, one-tailed) and reliably item-specific based on a randomization analysis (*p* = 0.02). Brain image shows segmented ROIs for one subject overlaid on their high-resolution T2w anatomical image. *n* = 23 in each group; \**p*< 0.05.

We then ran post-hoc tests to investigate the differentiation finding in right CA2/3/DG. Within the REM group, the neural differentiation score in right CA2/3/DG was significantly negative (t_22_ = –2.44, p = 0.01, d = –0.51, one-tailed), indicating that the A and B items making up violation condition pairs became less similar to each other than nonviolation condition pairmates. As illustrated in Figure S3, which plots the violation and nonviolation conditions separately, the greater differentiation effect in the REM group appears to be driven by nonviolation pattern similarity values being higher in the REM group than the other groups, rather than violation pattern similarity values being lower in the REM group; indeed, an exploratory contrast revealed that nonviolation pattern similarity values were significantly higher with REM than without REM (t_66_ = 2.12, p = 0.04, d = 0.52). This result suggests that, in the REM group, there may have been some integration of A and B items in the nonviolation condition that did not occur in the violation condition.

Given the significant effect of REM sleep in right CA2/3/DG, we next asked if this effect was item-specific. In other words, does B become more distinct from its specific A pairmate, not just generally more distinct from other items? We employed a randomization analysis where we shuffled the pair assignments of A and B 1000 times within each task condition and recalculated the neural differentiation score. If differentiation is item-specific, the actual neural differentiation score should be more negative (negative values indicate violation-related differentiation) than the shuffled distribution. Within each subject, we computed the z-score of the observed neural differentiation score relative to the mean and SD of 1000 shuffled scores, and tested the reliability of these z-scores across participants with a one-sample *t*-test against zero. This confirmed that violation-related neural differentiation in right CA2/3/DG in the REM group was item-specific (t_22_ = –2.62, p = 0.02, d = –0.54).

We did not have a specific prediction about whether the NREM and Wake groups would differ from each other. NREM sleep alone could yield some marginal benefit for neural differentiation compared with time spent awake. However, contrasting those two groups (contrast weights: REM: 0, NREM: −1, Wake: 1) revealed no difference in any subfield (left CA2/3/DG: t_66_ = –1.22, p = 0.23, d = –0.30; right CA2/3/DG: t_66_ = –0.22, p = 0.83, d = –0.05; bilateral CA2/3/DG: t_66_ = –0.88, p = 0.38, d = –0.22; left CA1: t_66_ = –0.30, p = 0.76, d = –0.07; right CA1: t_66_ = –0.52, p = 0.60, d = –0.13; bilateral CA1: t_66_ = –0.47, p = 0.64, d = –0.12).

Finally, we ran an exploratory analysis separating the differentiation scores in CA2/3 and DG subfields (Figure S4; raw pattern similarity values for each task condition can be found in Figure S5). The contrast testing the effect of REM versus NREM/Wake showed the predicted pattern in right DG only (right DG: t_66_ = 2.27, p = 0.03, d = 0.56, not significant after correcting for multiple comparisons; left DG: t_66_ = –0.37, p = 0.72, d = –0.09; bilateral DG: t_66_ = 1.03, p = 0.31, d = 0.25; left CA2/3: t_66_ = 0.45, p = 0.66, d = 0.11; right CA2/3: t_66_ = –0.14, p = 0.89, d = –0.03; bilateral CA2/3: t_66_ = 0.37, p = 0.71, d = 0.09), suggesting that violation-related neural differentiation in the REM group was primarily driven by the right DG. A post-hoc test showed that the neural differentiation score in the REM group in right DG was significantly negative (t_22_ = –2.31, p = 0.015, d = –0.48, one-tailed), and a randomization analysis further confirmed that this violation-related neural differentiation was item-specific (t_22_ = –2.48, p = 0.02, d = –0.52). Clearer differentiation in DG than CA2/3 is consistent with prior findings from the small number of studies to isolate DG (Wammes et al., 2022). Mirroring the result from the combined CA2/3/DG ROI, an exploratory contrast revealed that nonviolation pattern similarity values in right DG were significantly higher with REM than without REM (t_66_ = 2.77, p = 0.007, d = 0.68), suggesting there may have been some integration of A and B items in the nonviolation condition.

Our preferred explanation for why post-learning B snapshots become less similar to pre-learning A snapshots is that the representation of B moves away from A (relinquishing shared features and acquiring new features). An alternative explanation for the differentiation effects we observed is that the post-learning B snapshots include additional “noise” from the X and Y faces (Greve et al., 2018). According to this account, during violation events, when a participant sees A followed by X or Y while simultaneously reactivating B, the X and Y faces are bound to (or integrated with) both A and B, resulting in decreased similarity between pre-learning A (which does not incorporate this noise from X and Y) and post-learning B (which does). If true, then we might expect a negative relationship between preA/postB pattern similarity and a measure of B-X-Y integration. That is, the more that X-Y are integrated into B, the lower preA/postB pattern similarity should be. To measure B-X-Y integration, we calculated the Pearson correlation between the post-learning snapshot of B and the post-learning snapshots of X and Y. We then correlated this B-X-Y integration score with preA/postB pattern similarity across pairs within subject. This correlation was not reliably different from zero across subjects in the REM group in right CA2/3/DG (t_22_ = 0.93, p = 0.36, d = 0.19), and thus, X-Y “noise” does not appear to be driving the observed violation-related neural differentiation effect.

### Relationship between prediction and differentiation

As part of the hypothesized differentiation mechanism, we predicted that the degree to which A and B items differentiated would be related to the amount of B activation during violation trials (as observed by Kim et al., 2017). That is, after three instances of B following A in the stimulus sequence, how much did B come to mind when a participant was presented with A followed by a face? To measure this “B prediction” on violation trials (two violation trials per pair, with a unique face presented during each violation, which we refer to as faces X and Y), we calculated the Pearson correlation between the pre-learning snapshot of B and the pattern of activity evoked by the X and Y violation events, then averaged these values, yielding one B prediction score per pair in the violation condition (Figure 1b). Across pairs, we computed the correlation of this prediction score with the preA/postB pattern similarity values within each subject. If greater B prediction is related to more neural differentiation, this should yield a negative correlation value (i.e., more B prediction, less preA/postB pattern similarity). We then analyzed these within-subject correlation values across subjects at the group level.

Again, we hypothesized that the prediction-differentiation relationship would be stronger in the REM group compared to the NREM and Wake groups, specifically in left CA2/3/DG. However, the planned contrast testing for a difference between the REM group and the two other groups (contrast weights: REM: −1, NREM: 0.5, Wake: 0.5) revealed no significant difference in left CA2/3/DG (t_66_ = 1.26, p = 0.21, d = 0.31), or any other ROI (right CA2/3/DG: t_66_ = 0.92, p = 0.36, d = 0.23; bilateral CA2/3/DG: t_66_ = 1.34, p = 0.18, d = 0.33; left CA1: t_66_ = 0.63, p = 0.53, d = 0.16; right CA1: t_66_ = 0.41, p = 0.68, d = 0.10; bilateral CA1: t_66_ = 0.59, p = 0.56, d = 0.14; Figure S6).

An exploratory analysis testing the same contrast in CA2/3 and DG separately showed the predicted pattern in bilateral DG (t_66_ = 2.04, p = 0.046, d = 0.50; Figure S7), with higher levels of B prediction associated with decreased A-B pattern similarity in the REM group (one-sample *t*-test, t_22_ = –2.06, p = 0.03, d = –0.43, one-tailed); neither of these tests remained significant following correction for multiple comparisons. The contrast between REM and Wake/NREM was not significant in the remaining ROIs (left CA2/3: t_66_ = 1.80, p = 0.08, d = 0.44; right CA2/3: t_66_ = –0.17, p = 0.86, d = –0.04; bilateral CA2/3: t_66_ = 0.44, p = 0.66, d = 0.11; left DG: t_66_ = 1.18, p = 0.24, d = 0.29; right DG: t_66_ = 1.32, p = 0.19, d = 0.32). Nonetheless, the REM group numerically showed the predicted negative relationship between B prediction and A-B pattern similarity in all of these ROIs; in addition to bilateral DG, the negative relationship in the REM group also passed traditional significance levels in left CA2/3 (t_22_ = –2.50, p = 0.01, d = –0.52, one-tailed). However, the group-level contrast in left CA2/3 did not show that the REM group was different than NREM/Wake, so we interpret this with caution.

### Relationship between sleep and differentiation

We hypothesized that the degree to which items differentiated would not only depend on the amount of prediction during violation trials, but also on the composition of the intervening sleep period. Specifically, we expected that greater amounts of REM sleep would be associated with more violation-related differentiation (numerically, this should yield a negative correlation between REM duration and the differentiation score). We found that, within the REM group, minutes of REM sleep was not significantly correlated with the neural differentiation score in the preregistered ROIs (left CA2/3/DG: r = –0.12, p = 0.59; right CA2/3/DG: r = 0.10, p = 0.66; bilateral CA2/3/DG: r = 0.01, p = 0.96; left CA1: r = 0.36, p = 0.10; right CA1: r = –0.02, p = 0.94; bilateral CA1: r = 0.20, p = 0.38), nor was REM duration correlated with neural differentiation in the exploratory DG ROI that showed a difference between REM and Wake/NREM (left DG: r = –0.30, p = 0.17; right DG: r = 0.14, p = 0.52; bilateral DG: r = –0.09, p = 0.70).

We also ran exploratory analyses exploring the relationship between other sleep variables (TST, minutes of Stage 1, Stage 2, Stage 3, spindle density) and the neural differentiation score in CA2/3/DG and CA1 in all nap participants (NREM and REM groups combined). Stage 3 spindle density was positively correlated with the differentiation score in left and bilateral CA1 (left: r = 0.36, p = 0.02; bilateral: r = 0.34, p = 0.03), indicating that greater spindle densities were associated with *less* violation-related differentiation in these regions, but these correlations were not significant after correcting for multiple comparisons. There were no other significant correlations (all remaining *p*s > 0.08, see Table S1). Given the lack of significant correlations between sleep features and differentiation, we did not proceed with the additional multiple linear regression analyses proposed in the preregistration.

### Relationship between neural differentiation and behavioral reward learning

At the end of the main experiment, participants completed a secondary task with the aim of detecting a behavioral consequence of neural differentiation. We based our task on Wimmer and Shohamy (2012), who showed that reward values can spread across implicitly associated memories. Participants explicitly learned to associate either a reward or neutral outcome with the A scene from each pair; B scenes were never directly associated with the reward or neutral outcome during the learning phase. The rationale of the task is that, if the neural representations of A and B are more overlapping (i.e., less differentiated), then the learned reward associations should generalize more to the B scene pairmate. Therefore, we predicted *less* generalization to B scenes for pairs that experience *more* differentiation, specifically violation pairs in the REM group. However, we did not find any evidence of differences in generalization due to task condition or group assignment (see Supplementary Information for detailed methods and results).

## Discussion

Prior work from our lab found hippocampal differentiation when neural activity was measured approximately 24 hours after prediction violations (Kim et al., 2017). The current study provides a preregistered test of the hypothesis that REM sleep was a critical ingredient driving this representational change. In support of our hypothesis, we found greater differentiation in the REM group than the Wake and NREM groups in the same CA2/3/DG ROI as the previous study. Our effect was lateralized to the right, rather than the left, hemisphere, and was only significant at an uncorrected threshold. An exploratory analysis found that this effect was driven by right DG. We also hypothesized that the degree of neural differentiation would be correlated with the amount of prediction during violation events, and more so in the REM group than the Wake and NREM groups. This pattern was reliable at an uncorrected threshold in an exploratory bilateral DG ROI. Although our findings do not fully align with the preregistration, they are nevertheless consistent with the hypothesis that REM sleep is important for hippocampal neural differentiation. Indeed, all of the differentiation effects we obtained were larger in the REM group than the other two groups or apparent within the REM group alone, and were localized to the predicted region of the hippocampus (CA2/3/DG). More work is needed, but these results provide provisional evidence for our working hypothesis that neural dynamics during REM sleep allow spreading activation within the hippocampus to identify and coactivate memories marked for representational change (Norman et al., 2005). Below, we discuss notable aspects of the present work that can guide future research.

Our study builds on prior work that has examined representational change across periods of consolidation (i.e., time, including time spent asleep). Tompary and Davachi (2017) measured neural pattern similarity immediately post-learning and after one week. After a week of consolidation (but not immediately after encoding), unique objects associated with the same scene became integrated in medial prefrontal cortex (mPFC) and posterior hippocampus relative to objects associated with different scenes (see Ezzyat et al., 2018 for an example of consolidation-related differentiation in mPFC). Within the hippocampus, another study found that representations for object-word pairs that were studied at the same time (i.e., within a list) and subsequently remembered became more differentiated in the anterior compared to the posterior hippocampus (Cowan et al., 2021); this reorganization of memory representations along the long axis of the hippocampus only emerged after an overnight delay (also see Dandolo and Schwabe, 2018). Although sleep may have contributed to the effects reported in these studies, sleep was not included as an experimental variable. Cowan and colleagues (2020) reported correlational evidence linking sleep to the restructuring of memory representations, finding that fast spindle density during overnight sleep was associated with greater pattern similarity in ventromedial prefrontal cortex (vmPFC) for object-word pairs learned prior to sleep, and that this relationship was mediated by anterior hippocampal-vmPFC functional connectivity. Here, we manipulated the content of sleep (NREM only vs. combined NREM and REM) and included a quiet wake control group; this design allowed us to isolate the role of REM sleep. Interestingly, violation-related neural differentiation in the REM group appears to have been driven by increased (relative to NREM and Wake) pattern similarity in the nonviolation condition. These results are compatible with an interpretation where integration occurs — and is facilitated by REM sleep — in the nonviolation condition, but violation events prevent this integration from taking place. Our results thus contribute to an emerging picture in which learning demands determine if memories should be integrated or differentiated, and REM sleep drives those representational changes during a period of consolidation (Sterpenich et al., 2014).

The idea that REM sleep (as opposed to NREM sleep alone) was necessary for hippocampal differentiation appears to be at odds with other results showing that NREM sleep is the key sleep stage for hippocampal memory consolidation (Rasch et al., 2007; Marshall et al., 2006; Mednick et al., 2013). One reason why it might be challenging to appreciate the effects of REM sleep on memory consolidation is that it may not cause a simple increase or decrease in memory strength. Rather, as shown here, offline learning during REM may be important for shaping how memory representations relate to one another, either by pushing memories closer together or pulling them apart. The behavioral measures typically used to quantify declarative memory consolidation (e.g., memory for paired associates) may not be sensitive enough to detect these representational changes from REM sleep. Even in the current study, we did not find group-level differences in performance on our post-sleep behavioral measure (the reward learning task). Other studies that found a behavioral effect of REM sleep used measures sensitive to the structure of memory representations, such as interference between competing memories (Baran et al., 2010; McDevitt et al., 2015) or multi-item integration (Cai et al., 2009; Batterink et al., 2014; Abdou et al., 2024). Our study highlights how experiments can be designed to target the aspects of memory hypothesized to be REM-dependent.

Even though NREM sleep alone was not sufficient to bring about differentiation, it is still possible that NREM sleep contributed to the effects seen in the REM group. We did not test a REM-only condition, since it is not biologically normal to enter REM sleep without some amount of preceding NREM sleep — as such, we cannot claim that REM alone is sufficient. Rather, the combination of the two in sequence (i.e., a full sleep cycle) may be critical for bringing about these representational changes, as suggested by the “sequential hypothesis” (Diekelmann and Born, 2010; Giuditta et al., 1995). An alternative explanation of our results is that the differentiation we observed in the REM group was due to this group having more N2 sleep compared to the NREM group. However, we did not find any evidence of dose-dependent effects; no sleep variable, including time spent in N2 sleep, was significantly correlated with differentiation, suggesting that — even if the NREM group had more N2 — it would not have contributed to more differentiation. Having some amount of NREM and REM sleep, rather than high absolute amounts of either, might be the critical factor. Future research, including research using computational models that can manipulate the amount and structure of sleep (e.g., Singh et al., 2022), could address these open questions.

Whereas in our study differentiation required offline time spent asleep, other studies have found neural differentiation during or immediately following learning (Wanjia et al., 2021; Schlichting et al., 2015). For example, Wanjia and colleagues (2021) demonstrated that hippocampal differentiation abruptly happened at the “inflection point” in the learning process when participants showed clear behavioral evidence of discriminating similar cues. Our working hypothesis is that, if competition is not resolved during wake (e.g., there is not enough training to reach the inflection point), then offline learning during sleep can supplement and pull the competing representations further apart. An interesting extension of Wanjia et al. (2021) would be to test if pairs that do not reach the inflection point during learning, and presumably have not yet differentiated, show hippocampal differentiation following REM sleep without additional training. Future research is also needed to establish whether sleep is necessary to (1) stabilize already-differentiated hippocampal representations and (2) differentiate representations in long-term neocortical storage sites so that the behavioral memory benefits of differentiation (e.g., reduced interference, Favila et al., 2016) are upheld over the long-term (Ezzyat et al., 2018; Favila and Aly, 2023).

Although we obtained support for our hypothesis that REM sleep contributes to differentiation in CA2/3/DG (and DG alone), our effect was found in the right hemisphere instead of the left hemisphere, which was our main, preregistered ROI. Arguably, we erred in our preregistration by over-fitting to a single prior study (Kim et al., 2017). Reviewing the literature more broadly, there is strong support for differentiation in CA2/3/DG (or DG alone), and more than in CA1 (e.g., Wanjia et al., 2021; Wammes et al., 2022; Schapiro et al., 2012; Molitor et al., 2021). However, there is less support for differentiation being localized to the left hemisphere (for another example, see Bein and Davachi, 2024). In fact, many studies did not separate or analyze their results by hemisphere (Wanjia et al., 2021; Wammes et al., 2022; Dimsdale-Zucker et al., 2018; Molitor et al., 2021); Schapiro et al. (2012) did show a stronger numerical pattern of differentiation in right versus left CA2/3/DG. In retrospect, it would have been wise to treat results from these additional studies as a “prior” when filing the preregistration — with this hindsight, we might have been more agnostic about hemisphere and predicted an effect in CA2/3/DG with bilateral, left, and right ROIs.

Apart from these shortcomings in the preregistration, there were also some important differences between the present study and Kim et al. (2017) that may have contributed to discrepancies in the results. Kim et al. (2017) included a full night of sleep between prediction violations and the final measurement of representational change, whereas our study used a nap that was approximately 90 minutes in duration in the REM group. Using a daytime nap as our sleep intervention is both a strength and limitation, allowing us to: (1) minimize time-of-day effects, (2) implement a strong waking control without stressful sleep deprivation, and (3) experimentally manipulate the absence/presence of REM sleep (Mednick et al., 2003; McDevitt et al., 2015). However, a daytime nap is a shorter period of sleep, and it is possible that more representational change would occur over multiple sleep cycles in a full night. Another limitation is that we did not reach our target sample size of *N* = 102 (34 participants per group) because of pandemic-related data collection interruptions. Our final sample size of *N* = 69 (23 participants per group) may be underpowered to detect the full range of effects (for reference, Kim et al., 2017 reported their effects in one group of 32 participants). In particular, the REM group showed the expected direction of the prediction-differentiation relationship in right CA2/3/DG and right DG (the two ROIs that showed overall differentiation in the REM group), but the difference between groups was not statistically significant.

In conclusion, our findings provide suggestive evidence for how memory processing during REM sleep can complement the plasticity processes initiated during waking experience; when memories are identified as targets for representational change (here, as a result of prediction errors that occur during wake), some of the learning required to implement these changes may occur later, during REM sleep. These results provide initial, converging support for a link between REM sleep and representational change of individual memories in the hippocampus.

## Methods

### Preregistration

Study procedures, planned analyses, and predictions were preregistered (https://osf.io/p953t). We originally planned to test 34 participants in each of three experimental groups (total *N* = 102) as stated in the preregistration; this was based on an *a priori* power analysis. However, data collection was interrupted by the COVID-19 pandemic, and we made the decision to permanently stop data collection so that we could begin data analysis while in-person human subjects research was suspended.

### Participants

We collected complete datasets from 74 healthy, non-smoking adults (42 females, 8 left-handed, mean age = 20.4 years, range = 18-35 years) from the Princeton University community to participate in exchange for monetary compensation ($20 per hour). All procedures were approved by the Institutional Review Board for Human Subjects at Princeton University. All participants provided informed consent.

Following our preregistered criteria, two participants were excluded due to poor performance (2.5 SD below the mean) on the session 1 subcategory judgment task, and one participant was excluded due to poor performance (2.5 SD below the mean) on the session 2 reward learning task. Two participants did not agree to share their data publicly and were not included in our analysis. Our final sample included 69 participants (*n* = 23 per experimental group).

Participants reported normal or corrected-to-normal vision, with no history of neurological disorders, psychiatric disorders, major medical issues, or use of medication known to interfere with sleep. Participants also reported normally obtaining 6-9 hours of sleep per night (with weekday bedtime no later than 2am and wake time no later than 10am). The Epworth Sleepiness Scale (ESS; Johns, 1992) and the reduced Morningness-Eveningness Questionnare (rMEQ; Adan and Almirall, 1991) were used to screen for excessive daytime sleepiness (ESS score > 10) and extreme chronotypes (rMEQ < 8 or > 21). Heavy caffeine users (> 3 servings per day) were not enrolled in the study.

Participants were instructed to follow their regular sleep/wake schedule for one week prior to their study, and to spend at least 8 hours in bed the night prior to the study. To confirm adequate sleep was obtained the night before the study, participants completed an online, time-stamped sleep diary; if less than 6.5 hours of sleep was reported, participants were not tested and rescheduled for another day. Participants were asked to abstain from caffeine and alcohol starting at noon the day prior to the study.

### Stimuli

The stimulus materials were the same as in Kim et al. (2017) and consisted of color photographs of indoor and outdoor scenes, male and female faces, and natural and manmade objects presented on a gray background. Stimuli were projected on a screen behind the scanner and viewed with a mirror on the headcoil. Stimuli were presented using the Psychophysics Toolbox for MATLAB (http://psychtoolbox.org).

### Experimental procedures

#### Pre-study orientation and fMRI scan

Approximately 1-14 days prior to the scheduled study day, participants came to the lab for a study orientation appointment and pre-study fMRI scan. During this appointment, they were informed of all study procedures, provided informed consent, and completed study paperwork.

Participants also completed two runs of a functional localizer task in the scanner. Each run consisted of 15 blocks, with five blocks from each of three categories: faces, scenes, and objects. Participants categorized faces as male or female, scenes as indoor or outdoor, and objects as manmade or natural. Each stimulus was presented for 500 ms, followed by a blank interval of 1000 ms. Each block was 10 trials, and each 15 s block was followed by 15 s of fixation (i.e., rest). One run lasted approximately 7.8 minutes.

This scan served three main purposes: 1) to help participants become acclimated to the scanner environment prior to the full study day; 2) to help participants learn the button-press response mappings that were also used in the task on the full study day; and 3) to obtain data for training category-specific classifiers to potentially be used in our analyses (however, we do not report results using these data).

#### fMRI session 1

Session 1 began at 10:00 AM. Participants entered the MRI scanner and completed six runs of an incidental encoding task. Scenes and faces were presented one at a time, and participants performed a subcategory judgment task (“Is the scene indoor or outdoor?” “Is the face male or female?”). Participants fixated on a black central dot that changed to white when a response was recorded. Each trial began with a blink of the fixation dot to signal an upcoming stimulus, followed by the stimulus for 1000 ms, and a blank inter-stimulus-interval of 2000 ms. Each run consisted of 192 trials and lasted approximately 10 minutes.

The sequence of images followed an A-B pair structure: Each pair had scene A as the first item and a different scene B as the second item. These A-B pairs were inserted in the stream continuously amongst the other pairs, and participants were not made aware of this pair structure. Within each run, there were eight unique pairs for each of two task conditions (violation and nonviolation, see below), for a total of 16 pairs per run (16 pairs x 6 runs = 96 pairs total, 48 pairs per task condition). Scenes assigned to each pair and condition were randomized for each participant. Pair presentation order was also randomized for each participant with the constraint that the minimum and maximum distance between repetitions of the same pair was 2 and 20 pairs, respectively.

At the beginning of each run, the A and B members of each pair were shown once separately (i.e., B did not follow A) and randomly intermixed with scenes from other pairs. This first presentation of each scene is used to estimate each scene’s neural representation (“pre-learning snapshot”), uncontaminated by its pairmate.

Next, the scenes were shown together as a pair three times, interleaved with repetitions of other pairs. Each pair was assigned to one of two within-subject task conditions: violation and nonviolation. For pairs assigned to the violation condition, after the initial three repetitions of the pair, there were two “violation” events in which B failed to follow A. Instead, A was followed once by X and once by Y, where X and Y were novel faces. Following each violation event, the B item was subsequently “restudied” on its own, meaning it appeared in a novel context, not preceded by its pairmate A. Across the entire learning phase, violation pairs followed the sequence AB-AB-AB-AX-B-AY-B (intermixed with other pairs). Pairs in the nonviolation condition also had three initial repetitions, but no violation events. In order to match the frequency of B item exposures in the two conditions, the nonviolation condition also included two B “restudy” events. Across the entire learning phase, nonviolation pairs followed the sequence AB-AB-AB-B-B (intermixed with other pairs).

#### Offline period with EEG

Between fMRI sessions 1 and 2, all participants experienced between 90-120 minutes of controlled, offline time. Participants were randomly assigned to one of three offline conditions: wake, a 50-minute nap, or a 90-minute nap. Given that shorter naps tend to have less REM sleep than longer naps, these nap durations were chosen to increase the likelihood of having naps with and without REM sleep (Mednick et al., 2003; McDevitt et al., 2014, 2015; Schapiro et al., 2017a). For eventual data analysis, the naps were scored for sleep stages by an expert scorer according to standard criteria (Berry et al., 2012), and participants were re-grouped based on the content of their nap: NREM only naps (referred to as the NREM group) or naps with both NREM and REM (referred to as the REM group).

All participants returned to the lab at 12:45 PM and had electrodes attached for EEG recording. Participants in the wake condition experienced a period of “quiet wakefulness”; they sat in a chair in the EEG recording room and listened to podcasts for 90 minutes from approximately 1:30 PM - 3:00 PM. This condition was chosen to control for the reduced visual input and motor movement experienced during a nap, while still engaging participants enough to avoid them easily falling asleep. During this time, an experimenter monitored participants via EEG and a camera to make sure they remained awake, and alerted participants at the first sign of Stage 1 sleep.

Participants in the two nap conditions were given a nap opportunity beginning at approximately 1:30 PM. Sleep was monitored and quantified in real-time by an experimenter. Participants in the 50-minute nap condition were woken after 50 minutes of total sleep time was obtained, or at the first sign of REM sleep, whichever occurred first. The remaining amount of controlled offline time (up to 90 minutes) was filled with podcast listening. Participants in the 90-minute nap condition were woken after 90 minutes of total sleep time was obtained, but no later than 120 minutes after the beginning of their nap opportunity. For both nap conditions, the nap opportunity was ended if participants spent more than 30 consecutive minutes awake, and the remaining amount of offline time (up to 90 minutes) was filled with podcast listening.

Since it was expected that not all participants would fall asleep easily or stay asleep for the entire duration of their nap, we preregistered a contingency plan for how to adjust on the fly under very specific circumstances. We employed this adjustment procedure in 9 of the 69 participants included in our analyses. If a participant in the 50-minute nap condition had REM-onset sleep (i.e., REM precedes slow wave sleep), we did not wake the participant at the first sign of REM sleep. Instead, we let this participant sleep for up to 90 minutes, and analyzed their data as part of the REM group (*n* = 3). If a participant in the 90-minute nap condition did not obtain REM sleep, this dataset was analyzed as part of the NREM group (*n* = 2). If a participant in either nap condition did not fall asleep, or only had short, fragmented bouts of Stages 1 and 2 within the first 30 minutes of the nap period, the nap opportunity was ended and the participant listened to a podcast for the remaining amount of time. This dataset was analyzed as part of the Wake group (*n* = 4).

#### fMRI session 2

Session 2 began at 5:00 PM. Participants re-entered the MRI scanner and completed three separate tasks in the following order: 1) one run of post-learning B scene snapshots; 2) the reward association task (see Supplementary Methods for task details); and 3) one run of post-learning X and Y face snapshots. The post-learning snapshots followed the same procedure as the session 1 task (1000 ms stimulus duration, 2000 ms inter-stimulus-interval). For scene snapshots, all B scenes were shown again, in a random order, and participants made indoor/outdoor judgments. For face snapshots, all X and Y faces (used during the session 1 violation events) were shown in a random order, and participants made male/female judgments. Each snapshot run consisted of 96 trials and lasted approximately 5.3 minutes.

Throughout the study day, when participants were not being scanned or participating in the offline EEG session, participants were able to leave the lab and carry out their normal daily activities, but were instructed not to nap, consume caffeine, or exercise during this time. These breaks occurred from approximately 11:45 AM to 12:45 PM and 3:30 PM to 4:45 PM.

### fMRI data

#### fMRI data acquisition

MRI data were acquired on a 3T Siemens Skyra scanner using a 64-channel head coil at the Princeton Neuroscience Institute’s Scully Center for the Neuroscience of Mind and Behavior. Functional scans used a T2*-weighted multiband echo-planar imaging (EPI) sequence (TR=1500 ms, TE=40 ms, voxel size=1.5mm isotropic, flip angle=64*^◦^*, multiband factor=6, 72 slices manually aligned to top-of-AC, bottom-of-PC). These slices comprised a partial volume fully covering the occipital and temporal lobes. For fieldmap correction, two spin-echo field map volumes (TR=10330 ms, TE=68 ms) were acquired in opposite phase encoding directions. We collected the following anatomical scans: three whole-brain T1-weighted MPRAGE images (one collected during each fMRI scan session; TR=2300 ms, TE=2.98 ms, voxel size=1mm isotropic, flip angle=9*^◦^*, 176 slices, GRAPPA acceleration factor=2), one T2-weighted turbo spin-echo (TSE) image (acquired at the end of fMRI session 1; TR=11390 ms; TE=90 ms, voxel size=0.44 x 0.44 x 1.5 mm, flip angle=150*^◦^*, 54 slices acquired perpendicular to the long axis of the hippocampus, distance factor=20%), and three coplanar T1 FLASH images (one acquired during each fMRI scan session), but the FLASH images were ultimately not used in our preprocessing or analysis pipeline.

#### fMRI data preprocessing

Data were preprocessed using *fMRIPrep* 1.2.3 (Esteban et al., 2019, 2018), which is based on *Nipype* 1.1.6-dev (Gorgolewski et al., 2011, 2018). Many internal operations of *fMRIPrep* use *Nilearn* 0.4.2 (Abraham et al., 2014), mostly within the functional processing workflow.

##### Anatomical data preprocessing

A total of 3 T1-weighted (T1w) images were included within the input BIDS dataset (from the pre-study, session 1, and session 2 scans). All of them were corrected for intensity non-uniformity (INU) using N4BiasFieldCorrection (ANTs 2.2.0, Tustison et al., 2010). A T1w-reference map was computed after registration of 3 T1w images (after INU-correction) using mri_robust_template (FreeSurfer 6.0.1, Reuter et al., 2010). The T1w-reference was then skull-stripped using antsBrainExtraction.sh (ANTs 2.2.0), using OASIS as the target template. Brain surfaces were reconstructed using recon-all (FreeSurfer 6.0.1, Dale et al., 1999), and the brain mask estimated previously was refined with a custom variation of the method to reconcile ANTs-derived and FreeSurfer-derivedsegmentations of the cortical gray-matter of Mindboggle (Klein et al., 2017). Spatial normalization to the ICBM 152 Nonlinear Asymmetrical template version 2009c (RRID:SCR_008796, Fonov et al., 2009) was performed through nonlinear registration with antsRegistration (ANTs 2.2.0, Avants et al., 2008), using brain-extracted versions of both T1w volume and template. Brain tissue segmentation of cerebrospinal fluid (CSF), white-matter (WM) and gray-matter (GM) was performed on the brain-extracted T1w using fast (FSL 5.0.9, Zhang et al., 2001).

##### Functional data preprocessing

For each of the 18 BOLD runs per participant (across all tasks and sessions), the following preprocessing was performed. First, a reference volume and its skull-stripped version were generated using a custom methodology of *fMRIPrep*. A deformation field to correct for susceptibility distortions was estimated based on two echo-planar imaging (EPI) references with opposing phase-encoding directions, using 3dQwarp (AFNI 20160207, Cox and Hyde, 1997). Based on the estimated susceptibility distortion, an unwarped BOLD reference was calculated for a more accurate co-registration with the anatomical reference. The BOLD reference was then co-registered to the T1w reference using bbregister (FreeSurfer) which implements boundary-based registration (Greve and Fischl, 2009). Co-registration was configured with nine degrees of freedom to account for distortions remaining in the BOLD reference. Head-motion parameters with respect to the BOLD reference (transformation matrices, and six corresponding rotation and translation parameters) are estimated before any spatiotemporal filtering using mcflirt (FSL 5.0.9, Jenkinson et al., 2002). BOLD runs were slice-time corrected using 3dTshift from AFNI 20160207 (RRID:SCR_005927, Cox and Hyde, 1997). The BOLD time-series (including slice-timing correction when applied) were resampled onto their original, native space by applying a single, composite transform to correct for head-motion and susceptibility distortions. Gridded (volumetric) resamplings were performed using antsApplyTransforms (ANTs), configured with Lanczos interpolation to minimize the smoothing effects of other kernels (Lanczos, 1964).

After preprocessing with *fMRIPrep*, the first nine volumes and last five volumes of each functional scan were discarded. Then all functional scans were additionally high-pass filtered (1/128 Hz cutoff) and *z*-scored using *Nilearn* prior to further analysis.

##### Hippocampal segmentation

Hippocampal subfields were defined using the Automated Segmentation of Hippocampal Subfields (ASHS) toolbox (Yushkevich et al., 2015) and a database of manual MTL segmentations from a separate set of participants (Aly and Turk-Browne, 2016a,b). Each participant’s fMRIPrep-preprocessed T1w template and their raw T2w TSE image were submitted as input to ASHS. The resulting segmentations were used to make masks for the CA1, CA2/3, dentate gyrus (DG), and combined CA2/3/DG subfields in both hemispheres. These masks were then transformed and resampled to match the functional data.

#### Measuring neural differentiation

We followed the same fMRI data analysis procedure as Kim et al. (2017). The goal of this analysis was to measure how much the neural representation of B items moved away from the original representation of their A pairmate, and compare these pattern similarity values for violation and nonviolation pairs. We used the pre-learning snapshot of A as the baseline for representational change to avoid confounds due to item frequency that could be introduced by using the post-learning snapshot of A since A items in the violation condition were presented two more times than A items in the nonviolation condition.

For each pair, we computed the Pearson correlation between the pre-learning snapshot of A and post-learning snapshot of B. Pre- and post-learning snapshots were defined as the spatial pattern of activity elicited by each item in a particular ROI at the peak of the hemodynamic response (4.5 s after image onset). We transformed Pearson’s *r* to Fisher’s *z*, computed the average pattern similarity for pairs within the violation and nonviolation task conditions, and then calculated the difference of violation minus nonviolation conditions. We refer to this difference score as the neural differentiation score; negative values indicate decreased pattern similarity (i.e., less neural overlap or more differentiation) in the violation compared to nonviolation condition.

In order to test if neural differentiation effects are item-specific (i.e., does B become more distinct from A specifically, not just generally more distinct from other items?), we performed a randomization analysis. For each participant, we shuffled the pair assignments of A and B 1,000 times within each task condition. For each shuffle, we recalculated the average pattern similarity for violation and nonviolation (shuffled) pairs, and then computed the violation minus nonviolation neural differentiation score. If differentiation is item-specific, the original neural differentiation score should be more negative than the shuffled distribution. This was quantified by computing, for each participant, the z-score of the true neural differentiation score relative to the mean and SD of the null distribution of 1,000 shuffled differentiation scores.

#### Relating prediction to differentiation

The goal of this analysis is to examine how differentiation of A and B items relates to the amount of B activation during the two violation events, when scene A was followed by faces X and Y instead of the expected B pairmate. This analysis was only performed for the violation condition. To measure B prediction on violation trials, we calculated the Pearson correlation between the pre-learning snapshot of B and the pattern of activity evoked by the X and Y items. Specifically, for each pair, we correlated Bpre/X and Bpre/Y, then Fisher *z*-transformed the resulting correlation coefficients and averaged these two values to provide a single estimate of B prediction for each pair. Across pairs, we calculated the correlation of the B prediction score with the Apre/Bpost neural pattern similarity score, resulting in one Pearson’s *r* value for each subject, which was transformed to Fisher’s *z* for statistical analysis at the group level.

#### Measuring integration of X and Y faces with B scenes

We ran a control analysis to rule out the alternative explanation that post-learning B snapshots appear less similar to pre-learning A snapshots in the violation condition because the post-learning B snapshots include additional noise from faces X and Y (Greve et al., 2018). To measure the amount of B-X-Y integration for each B item in the violation condition, we calculated the Pearson correlation between the post-learning snapshot of B and the pattern of activity evoked by the corresponding X and Y faces during their post-learning snapshot phase. Specifically, we correlated Bpost/Xpost and Bpost/Ypost, then transformed these values to Fisher’s *z*, and averaged them to arrive at a single value of B-X-Y integration. Across all violation pairs within each subject, we correlated this B-X-Y integration score with the corresponding Apre/Bpost neural pattern similarity score, resulting in one Pearson’s *r* value for each subject, which was transformed to Fisher’s *z* for statistical analysis at the group level.

### EEG data processing

#### EEG data acquisition

Electroencephalogram (EEG) data were acquired using Ag/AgCl active electrodes (Biosemi Active Two) from 64 scalp EEG locations. We followed standard polysomnography procedures and additionally recorded data from two electrooculogram (EOG) locations (LOC and ROC), two submental electromyogram (EMG) locations, two electrocardiogram (ECG) locations, and two mastoids. Data were sampled at 512 Hz.

#### EEG data preprocessing and sleep scoring

Although we recorded EEG data during Wake sessions, those data were only used to ensure wakefulness and were not further analyzed. Nap EEG data were preprocessed using BrainVision Analyzer 2.0 (BrainProducts, Munich, Germany). For sleep scoring purposes, channels used for scoring (LOC, ROC, C3, C4, O1, O2) were re-referenced to the average signal of the left and right mastoid; in cases where one mastoid electrode became detached, a single mastoid was used (*n*=3). The data were bandpass filtered between 0.3 and 35 Hz with a 60 Hz notch filter and down-sampled to 256 Hz. One EMG channel was subtracted from the other to create one bipolar EMG channel. The EMG channel was bandpass filtered between 10 and 70 Hz. Data were visually scored for sleep stages in 30 s epochs following standard criteria (Berry et al., 2012) using the Hume 1.0.4 toolbox for Matlab (https://github.com/jsaletin/hume). Note that sleep records for two participants were unable to be scored for sleep staging due to an excessively noisy signal (*n*=1, NREM group) and the EEG amplifier battery dying in the middle of the nap session (*n*=1, REM group). These datasets were not included in analyses that depend on knowing the precise amount of time spent in sleep stages, but they were able to be included in group-level analyses since the experimenter was still able to determine if their nap contained REM sleep or not.

For further analysis of sleep EEG data, artifacts (large movements, arousals, and rare, large deflections in single channels) during sleep were visually identified and rejected in 5 s chunks. Problematic channels were interpolated. Sleep spindles were detected during Stage 2 and Stage 3 using a wavelet-based algorithm (Wamsley et al., 2012; Warby et al., 2014). Spindle densities were calculated by dividing the number of discrete spindle events by time spent in the corresponding sleep stage. We report spindle data from one exemplar centroparietal electrode (CPz).

### Statistical analyses

Data analysis was performed in RStudio using R 4.2.1. Accuracy on the behavioral cover task was not normally distributed. Therefore, we used the WRS2 package in R to perform robust ANOVA on 20% trimmed means (Mair and Wilcox, 2020). Specifically, we used the t1way function to test for overall group differences in accuracy, and the bwtrim function to compute a two-way mixed model ANOVA for analyses including both within- and between-subject factors.

We used independent samples *t*-tests to test for differences between the two nap groups on sleep architecture measures. When the data within either group were not normally distributed, we report a Wilcoxon rank sum test instead of the independent samples *t*-test. If the assumption of homogeneity of variances was violated, we report Welch’s *t*-test.

To directly test our main hypotheses, we ran planned contrasts to examine differences between groups. The first contrast compared the REM group to the two groups that did not have REM sleep (contrast weights REM: −1, NREM: 0.5, Wake: 0.5). Next, we further asked if the NREM group was different from the wake group (contrast weights REM: 0, NREM: −1, Wake: 1). Since we ran the planned contrasts in six individual ROIs, the Bonferroni adjusted alpha level for each set of tests was 0.05/6 = 0.008.

Contrasts that suggested the REM group was different from the Wake and NREM groups were followed up with one-sample *t*-tests against zero to test whether neural measures were significantly positive or negative in each group, with a Bonferonni adjusted alpha level of 0.05/3 = 0.017 (*k* = 3 for three groups). Since we preregistered the direction of the expected effects in the REM group, we report one-tailed *p*-values in the REM group (our preregistration did not specifically state we would use one-tailed tests but we did preregister the direction of effects).

For the randomization analysis, we computed the *z*-score of the true neural differentiation score relative to the mean and SD of the null distribution of 1,000 shuffled differentiation scores within each subject, and tested the reliability of these *z*-scores across participants with a one-sample *t*-test against zero.

We used a two-way repeated measures ANOVA to analyze the reward phase data with both task condition (violation/nonviolation) and repetition as within-subject repeated measures. We used mixed-model ANOVAs for analyses including both within- and between-subject factors; for example, to test for differences in the Decision score we included task condition (violation/nonviolation) as the within-factor and group (Wake/NREM/REM) as the between-factor. Pearson correlations examined the relationship between neural measures and sleep, and between neural measures and behavior.

## Data availability

The MRI, EEG, and behavioral data from this study will be deposited on Openneuro.org prior to publication.

## Code availability

The analysis scripts will be available on GitHub prior to publication.

## Author contributions

E.A.M., G.K., N.T.-B., and K.A.N. designed the study. E.A.M. collected and analyzed the data in consultation with K.A.N. E.A.M. drafted the manuscript and all authors edited the manuscript.

## Acknowledgments.

We thank Monika Schönauer and James Antony for helpful discussions and comments on an earlier version of this manuscript.

## Funding information

This work was supported by the National Institutes of Health (NIMH R01-MH06945 to K.A.N and N.T.-B. and NIMH K99-MH126154 to E.A.M.).

## Competing interests

The authors declare no competing interests.

## Supplementary Information

### Supplementary Methods

#### fMRI Session 2: Reward learning task

The reward association task was based on the task used by Wimmer and Shohamy (2012). Below we detail how the A scene from each pair was explicitly associated with either a reward or neutral outcome followed by a test of generalization of the learned reward value to the B scene pairmate. This differs from Wimmer and Shohamy (2012), who explicitly associated the reward/neutral outcomes with their “B” item and tested generalization to their “A” item.

Participants completed two cycles of the task; each cycle included three sub-tasks (one run of the familiarization task, two runs of the reward prediction task, and one run of the decision task). We split our scene pairs from session 1 into two separate sets; each set consisted of 48 scene pairs, with 24 pairs coming from each task condition (violation and nonviolation). The task proceeded in two separate cycles, with one set assigned to each cycle. A full cycle of the task was completed before beginning the second cycle.

During the familiarization task, participants were exposed to associated reward or neutral outcomes for each pair’s A scene; no B scenes appeared during this phase. Participants were explicitly instructed to try to learn whether each scene was associated with a reward or neutral outcome. Within each of the two original task conditions (violation and nonviolation), half of the A scenes were paired with a reward (A+), and the other half were paired with a neutral outcome (A-). Each A scene was presented once, followed by its associated reward or neutral outcome (either a picture of a $1 bill or a gray rectangle), in a random order. No responses were required during this phase. Each trial consisted of a scene shown for 1000 ms, followed by a blank 500 ms inter-stimulus-interval, and the outcome image for 1000 ms. The inter-trial-interval was 2000 ms. Each run consisted of 48 trials and lasted approximately 4 minutes.

During the reward prediction task, participants saw A scenes and were asked to predict if each scene led to a reward or neutral outcome; B scenes did not appear during this phase. Each A scene was tested two times in each run (four times total) in a pseudo-random order, such that all A scenes were tested once before any scene was tested a second time. After making their prediction, participants received feedback about the actual outcome, and were able to win or lose “bonus points” based on their prediction. Bonus points were assigned following these rules: If a participant predicted reward and the actual outcome was reward, they won +1 point. If a participant predicted reward and the actual outcome was neutral, they lost −1 point. If a participant predicted neutral, they did not win or lose any points, regardless of the actual outcome. This reward structure was aimed at optimizing the learning rate while keeping the false alarm rate low (e.g., net earning would be 0 if participants predicted ‘reward’ on all trials). Bonus points were converted to bonus monetary compensation at the end of the study. On each trial of the task, a scene image was shown alone for 2500 ms and participants made their prediction response within this timeframe. The scene remained on the screen as the following pieces of feedback were added in sequence: Participants saw a text version of their response (e.g., “You responded REWARD”) for 1000 ms, followed by the actual outcome image (a $1 bill or gray rectangle) for 1000 ms, followed by their winnings (either “You win +1” in green text, or “You lose −1” in red text, or “0” in black text if they did not win or lose points) for 1000 ms. The inter-trial-interval was 500 ms. Each run consisted of 96 trials and lasted approximately 10 minutes.

During the decision task, we assessed preference for A and B scenes. Each decision trial either pitted a rewarded A scene against a neutral A scene (A+ vs. A-), or a rewarded-by-association B scene against a neutral-by-association B scene (B+ vs. B-), and participants were tasked with choosing the scene they thought was more likely to lead to winning a reward. Since B scenes were not shown during the familiarization or reward prediction phases, the reward/neutral status of each B scene was only based on whether its A pairmate was rewarded or not rewarded. Performance was operationalized as the proportion of trials where the rewarded scene (A+ or B+) was chosen (we call this the ‘decision score’). We expected these scores to be very high for A+ vs. A-trials due to explicit learning of the A reward associations. Critically, if the learned A+ or A-reward associations generalized to their B scene pairmates, then there should be a preference for B+ scenes (i.e., they should choose the B+ at a rate higher than chance). This is the outcome we predict will be strongest for pairs whose neural representations are more overlapping (i.e., less differentiated). When analyzing the data, pairs that showed a greater decision bias for B+ than A+ (which was a rare occurrence, < 1% of pairs) were excluded from the analysis. Trials with no response were imputed to reflect approximately chance level performance across those no response trials within a participant, rather than removing no response trials from the analysis (i.e., if a participant had an even number of no response trials, half of those trials were randomly assigned an accuracy of 1 and the other half were assigned an accuracy of 0 for an average of 0.5 for those trials; if a participant had an odd number of no response trials, the “extra” trial was assigned an accuracy of 0). This procedure of imputing no response trials was included in our preregistration because we wanted all pairmates to have an equal number of trials represented in the analysis (i.e., we did not want to remove trials), and it has been argued that always scoring trials with a missing response as ‘incorrect’ does not accurately reflect memory failures, which at worst should produce 50/50 guessing (Potter et al., 2018). This imputation procedure was employed in 26 participants (range of number of trials with no response in these participants: 1-55); for these 26 participants, their overall accuracy after imputing no response trials was 72.98% vs. 72.34% if we had scored all no response trials as ‘incorrect’.

Each A and B scene appeared on four separate decision trials, each time pitted against a different scene counterpart (in other words, all stimulus pairings during the decision phase were unique). However, the pair shufflings were consistent for A and B trial types. For example, if A23+ was pitted against A16-on a trial, then B23+ was pitted against B16-on another trial. Further, the stimulus pairings for each decision trial always came from the same original task condition (violation or nonviolation), and were matched for indoor/outdoor subcategory. The screen location (left or right) of the rewarded image was counterbalanced across the four decision trials for each rewarded image. On each trial of the task, the two scene images were shown together for 2500 ms and participants were instructed to respond within this timeframe. The instructions emphasized that a response should be made on each and every trial, and it was better to guess than not respond at all. Upon a response, a blue frame surrounded the chosen scene image for the remainder of the trial. The inter-trial-interval was 500 ms. Participants earned a bonus point for correctly choosing the rewarded scene on each trial, which was converted to a monetary bonus at the end of the experiment, but feedback about accuracy was not given on a trial-by-trial basis. Each run consisted of 192 trials and lasted approximately 10 minutes.

## Supplementary Notes

### Supplementary Note 1: Additional preregistered analysis of neural pattern similarity

For analyzing neural pattern similarity, the omnibus test of our full design is a 2×3 mixed-model ANOVA with task condition as the within factor (violation, nonviolation) and group as the between factor (Wake, NREM, REM) (see Figure S3). In our preregistration, we wrote that we expected to find 1) a main effect of task condition, with lower pattern similarity in the violation compared to nonviolation condition, and 2) a significant condition x group interaction. In right CA2/3/DG, there was a main effect of task condition in the predicted direction (F_1,66_ = 4.17, p = 0.045), but the condition x group interaction was not significant (F_2,66_ = 2.42, p = 0.08). In left and bilateral CA1, there was a main effect of task condition where pattern similarity was greater in the violation than nonviolation condition (left CA1: F_1,66_ = 6.48, p = 0.01; bilateral CA1: F_1,66_ = 4.86, p = 0.03), but the condition x group interaction was not significant in either region (left CA1: F_2,66_ = 0.10, p = 0.90; bilateral CA1: F_2,66_ = 0.44, p = 0.65). There were no other significant ANOVA results, including no main effects of group.

### Supplementary Note 2: Reward learning task behavioral results

After an intial exposure trial, each A scene was tested four times across the reward learning phase. On each trial, participants saw an A scene and predicted if that scene was associated with a reward or neutral outcome. After making their prediction, participants received feedback about the actual outcome. Accuracy increased across the four learning repetitions (main effect of repetition, F_3,204_ = 215.44, p < 0.001) and was excellent by the fourth and final learning repetition (mean = 0.98, sd = 0.02), indicating that the explicit A scene reward associations were well-learned (Figure S8a). There was no interaction between pair type (violation, nonviolation) and learning repetition (F_3,204_ = 0.48, p = 0.70; Figure S8b), and no main effect of pair type (F_1,68_ = 0.52, p = 0.47).

During critical decision test trials, we either pitted a rewarded A scene against a neutral A scene (A+ vs. A-), or a rewarded-by-association B scene against a neutral-by-association B scene (B+ vs. B-). Since B scenes were not shown during the preceding explicit learning phase, the reward/neutral status of each B scene was only based on whether its A pairmate was rewarded or not rewarded. Participants were instructed to choose the scene they thought was more likely to lead to winning a reward. A decision score was computed as the proportion of trials where the rewarded scene (A+ or B+) was correctly chosen over the neutral scene. As expected, decision scores were very high for A trials (trials pitting A+ vs. A-; mean = 0.98, sd = 0.03; Figure S8c) and were near chance for B trials (trials pitting B+ vs. B-; mean = 0.50, sd = 0.05; Figure S8d). Mixed-model ANOVAs with task condition (violation, nonviolation) as the within-factor and group (Wake, NREM, REM) as the between-factor confirmed no significant main effects or interactions for either A trial or B trial decision scores (all *p*s > 0.20). We considered a decision score significantly greater than chance (0.50) on B trials as evidence of generalization, but did not observe that in any condition (all *p*s > 0.07).

We anticipated that it might be difficult to observe behavioral effects at the group level, and were interested in relating variance in this behavioral outcome to variance in neural differentiation. We computed the difference in the decision score for violation minus nonviolation pairs in each participant (Figure S8e); negative values indicate less generalization for violation compared to nonviolation pairs, in line with our hypothesis. We then correlated this behavioral measure with the neural differentiation score in right CA2/3/DG across participants. We predicted a positive relationship, such that more negative neural differentiation scores would be associated with more negative behavioral difference scores (i.e., more violation-related neural differentiation, less violation-related generalization). However, we did not observe any significant relationships between our neural and behavioral measures in all participants together (r = –0.0012, p = 0.99), or within the REM group specifically (r = –0.16, p = 0.47). As such, we did not proceed with the mediation analysis proposed in the preregistration.

## Supplementary Figures

**Figure S1.**
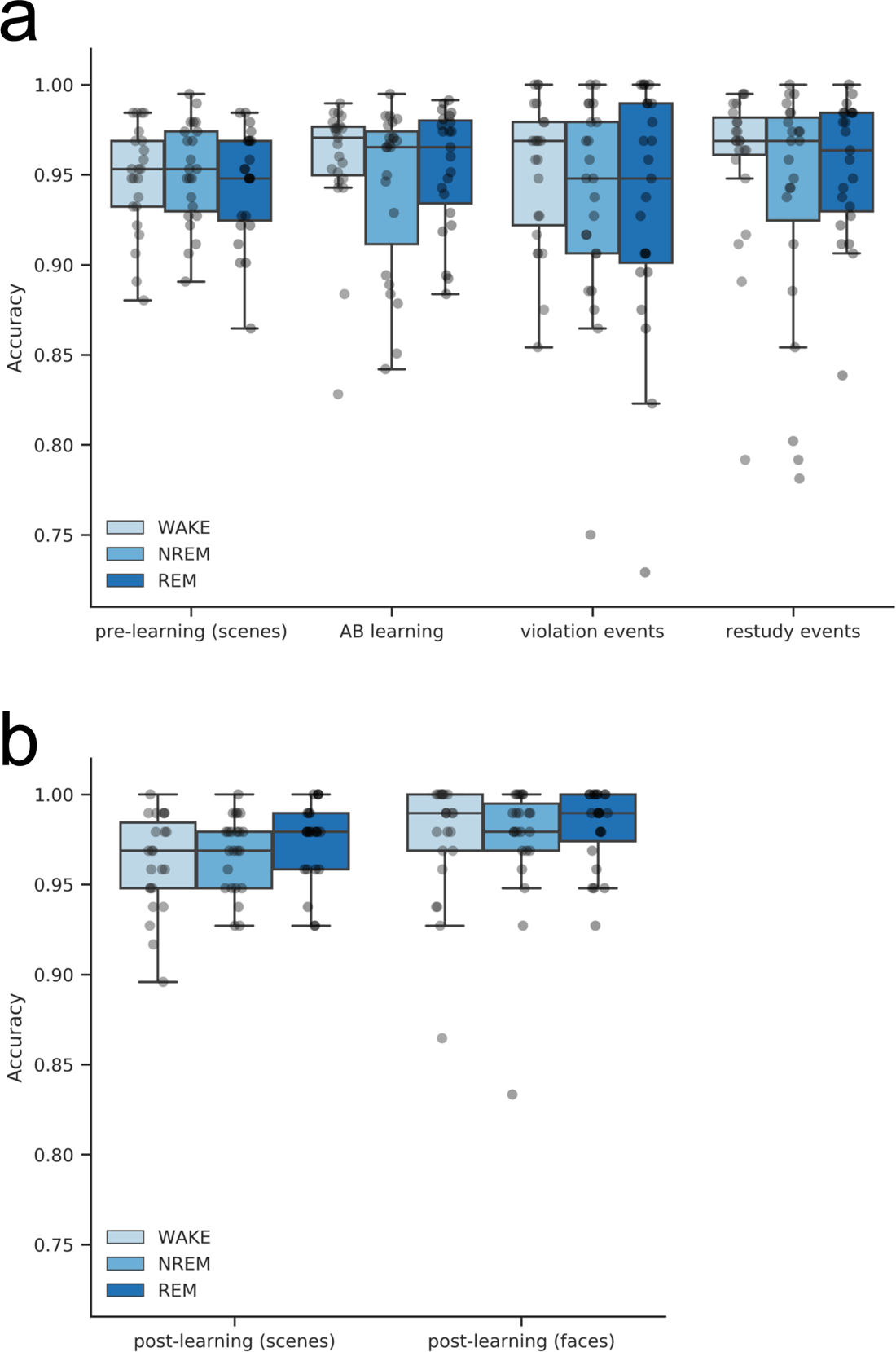
Behavioral performance. (A) Accuracy on the session 1 cover task for the following trial types: pre-learning scene snapshots, three repetitions of A-B learning, violation events (seeing faces X or Y instead of scene B), and restudy events (B not following A). (B) Accuracy on the session 2 cover task for the post-learning scene and face snapshots. n = 23 in each group.

**Figure S2.**
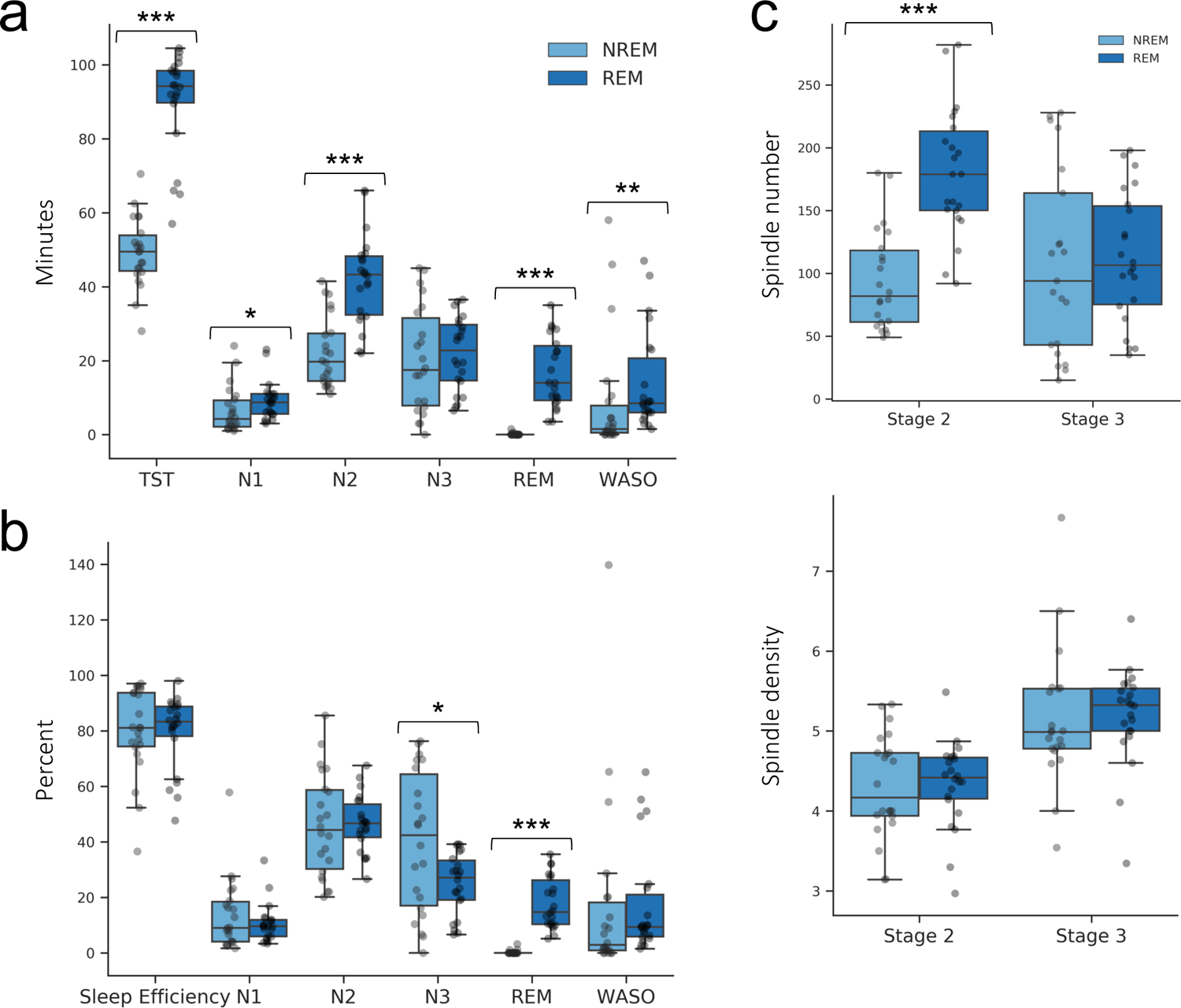
Nap sleep architecture. (**A**) Minutes of total sleep time (TST), Stage 1 (N1), Stage 2 (N2), Stage 3 (N3), REM sleep, and wake after sleep onset (WASO). (**B**) Sleep variables as a percentage of total sleep time. Sleep efficiency is total sleep time/time spent in bed trying to nap. (**C**) Number of spindle events (top panel) and spindle density (bottom panel) in Stage 2 and Stage 3 sleep, derived from channel CPz. *n* = 22 in each group; \**p*< 0.05, \*\**p*< 0.01, \*\*\**p*< 0.001.

**Figure S3.**
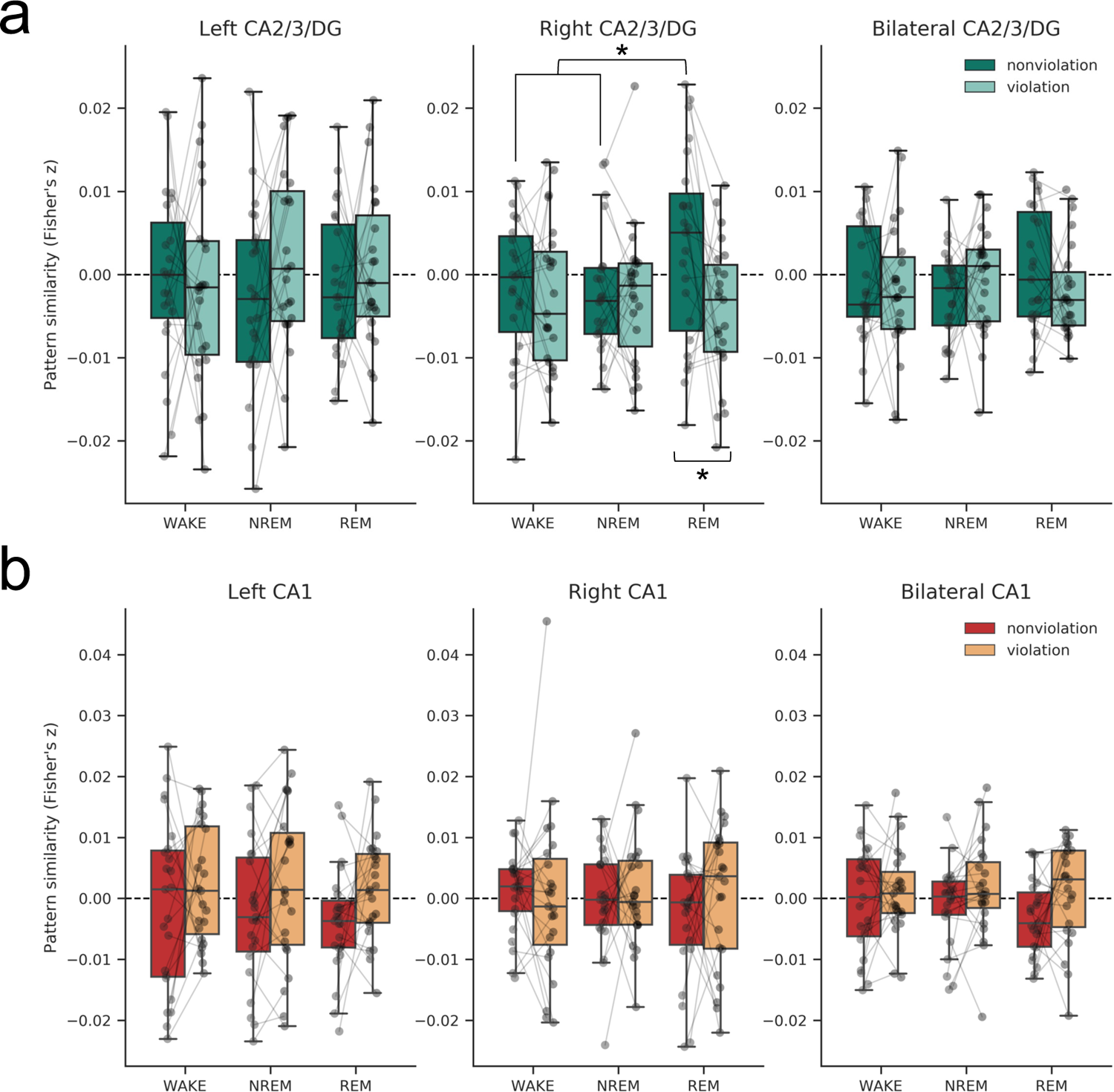
Pattern similarity by task condition in (a) CA2/3/DG and (b) CA1. The REM group showed the predicted pattern of results in right CA2/3/DG, with pattern similarity in the violation condition significantly less than in the nonviolation condition (*p* = 0.02, note this is the same finding depicted in Figure 2a in the main text). An exploratory contrast revealed that nonviolation condition pattern similarity values were significantly higher in the REM group than Wake/NREM groups (*p* = 0.04). *n* = 23 in each group; \**p*< 0.05.

**Figure S4.**
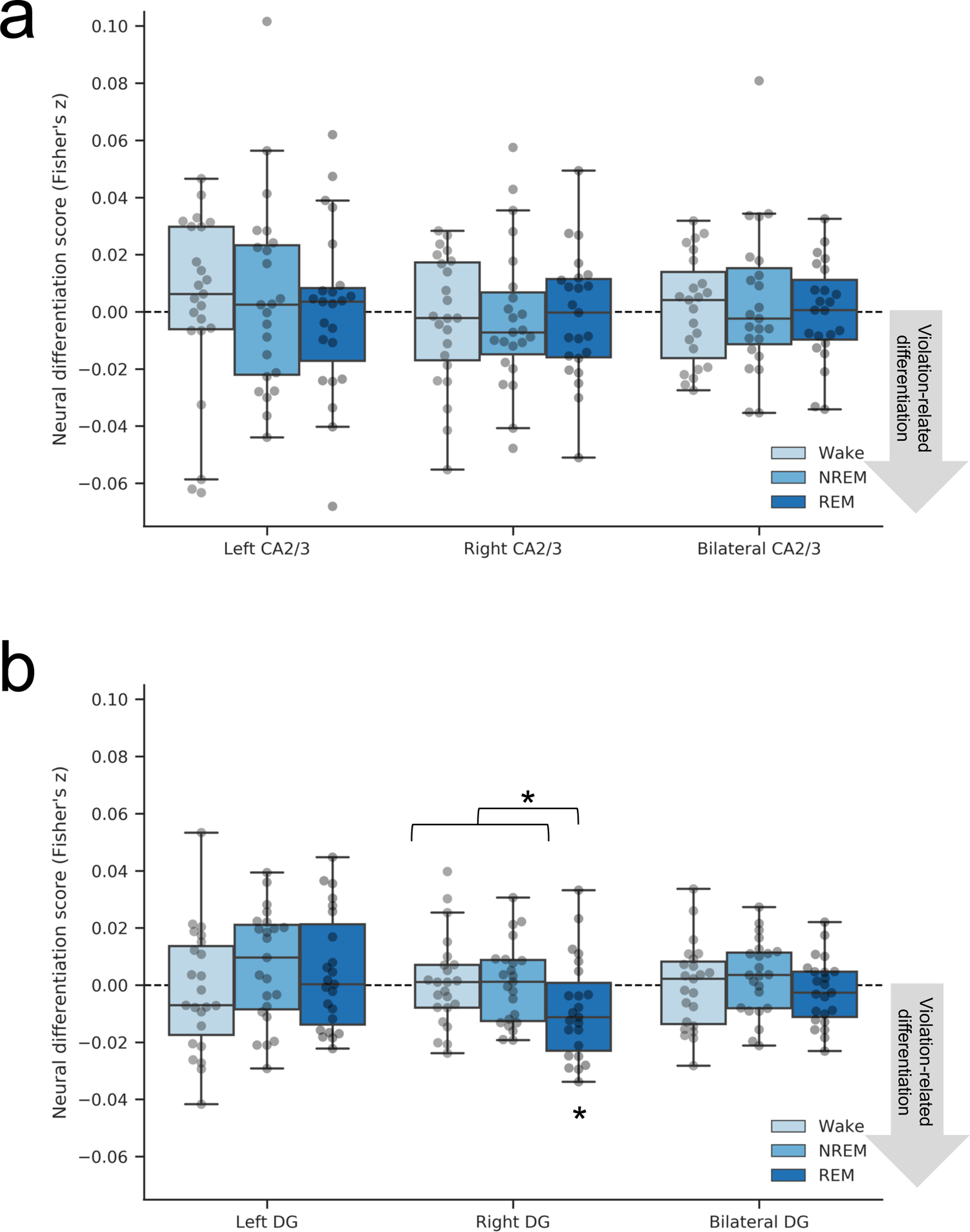
Neural differentiation in (a) CA2/3 and (b) DG separately. Neural differentiation scores are the difference in pattern similarity for the violation minus nonviolation task conditions. A contrast revealed more violation-related neural differentiation in the REM group compared to the Wake and NREM groups in right DG (*p* = 0.03, not significant following correction for multiple comparisons). Within the REM group, the neural differentiation score was significantly different from zero (*p* = 0.015, one-tailed) and reliably item-specific based on a randomization analysis (*p* = 0.02). *n* = 23 in each group; \**p*< 0.05.

**Figure S5.**
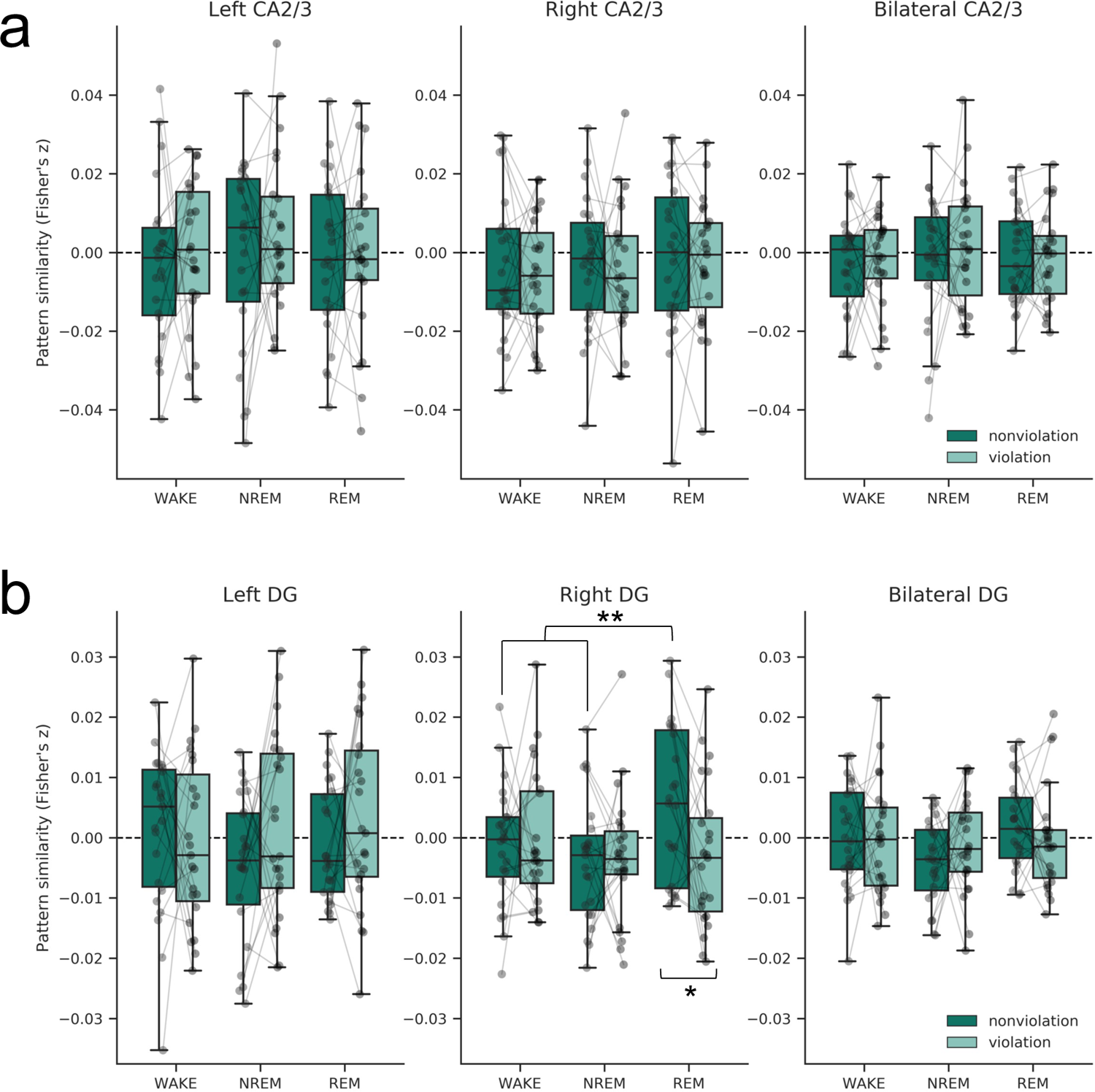
Pattern similarity by task condition in (a) CA2/3 and (b) DG separately. The REM group showed the predicted pattern of results in right DG, with pattern similarity in the violation condition less than in the nonviolation condition (*p* = 0.03, note this is the same finding depicted in Figure S4b). An exploratory contrast revealed that nonviolation condition pattern similarity values were significantly higher in the REM group than Wake/NREM groups (*p* = 0.007). *n* = 23 in each group; \**p*< 0.05, \*\**p*< 0.01.

**Figure S6.**
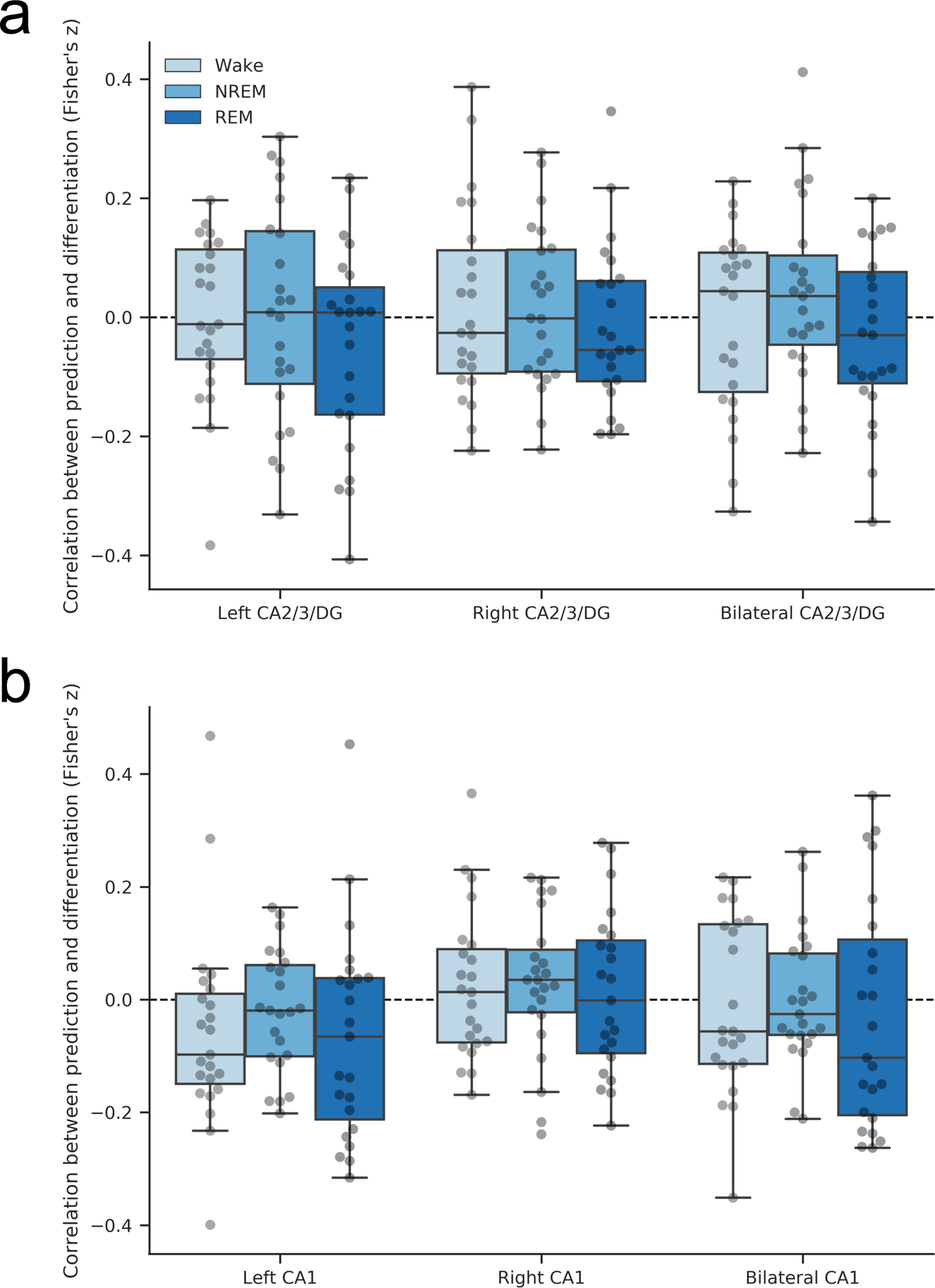
Relationship between prediction and differentiation in (a) CA2/3/DG and (b) CA1. We hypothesized that a negative correlation should exist between prediction and differentiation (i.e., higher levels of B activation during violation events should be associated with decreased A-B pattern similarity). However, there was no reliable relationship between B prediction and neural differentiation in any group in either CA2/3/DG or CA1 ROIs. *n* = 23 in each group.

**Figure S7.**
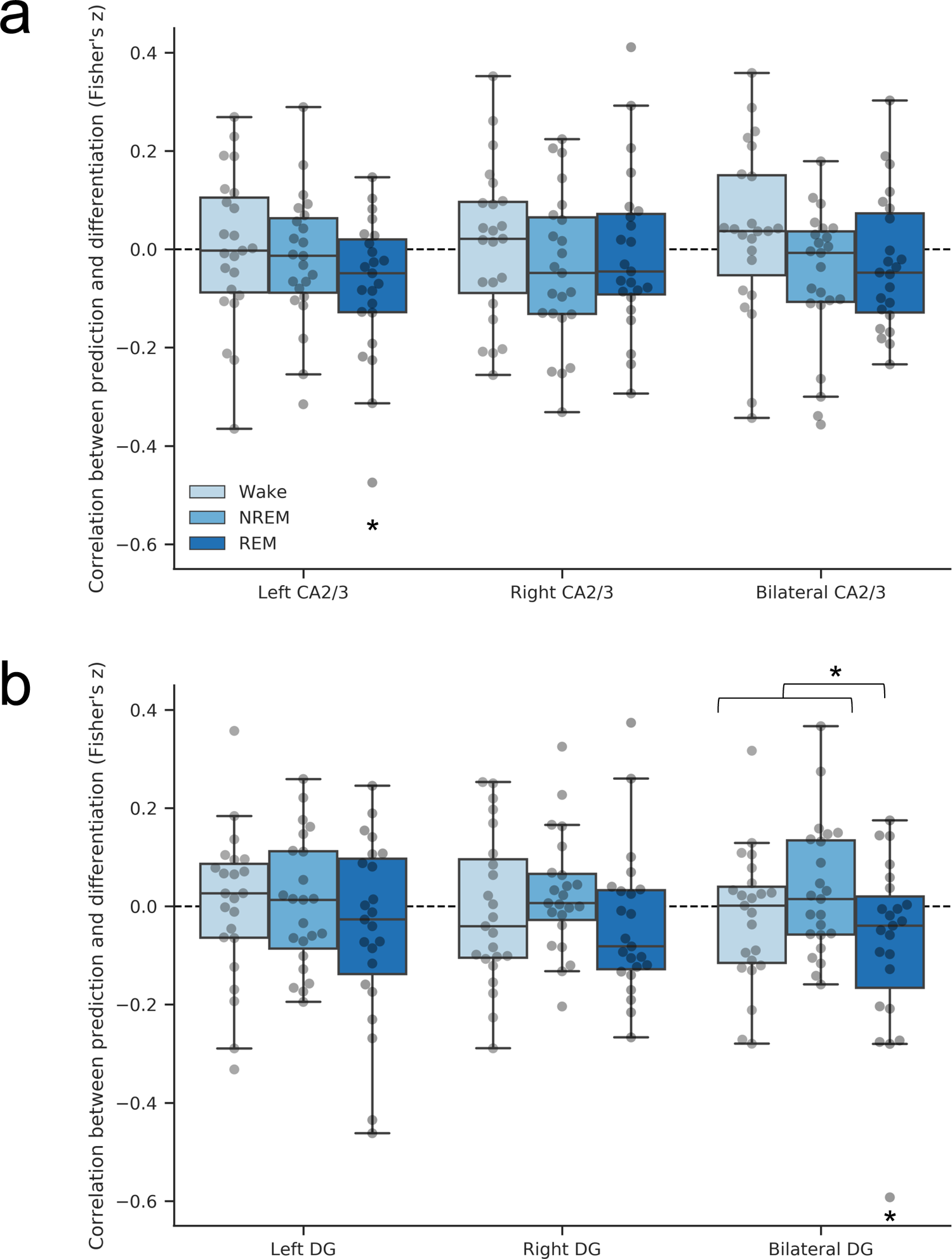
Relationship between prediction and differentiation in (a) CA2/3 and (b) DG separately. A contrast testing for a difference between the REM group and the two other groups showed a difference in bilateral DG (*p*=0.046, not significant after correcting for multiple comparisons), with the REM group showing the predicted negative relationship between B prediction and A-B pattern similarity (one-sample *t*-test, *p*=0.03, one-tailed). The relationship was also significantly different from zero in the REM group in left CA2/3 (*p* = 0.01, one-tailed). *n* = 23 in each group; \**p*< 0.05

**Figure S8.**
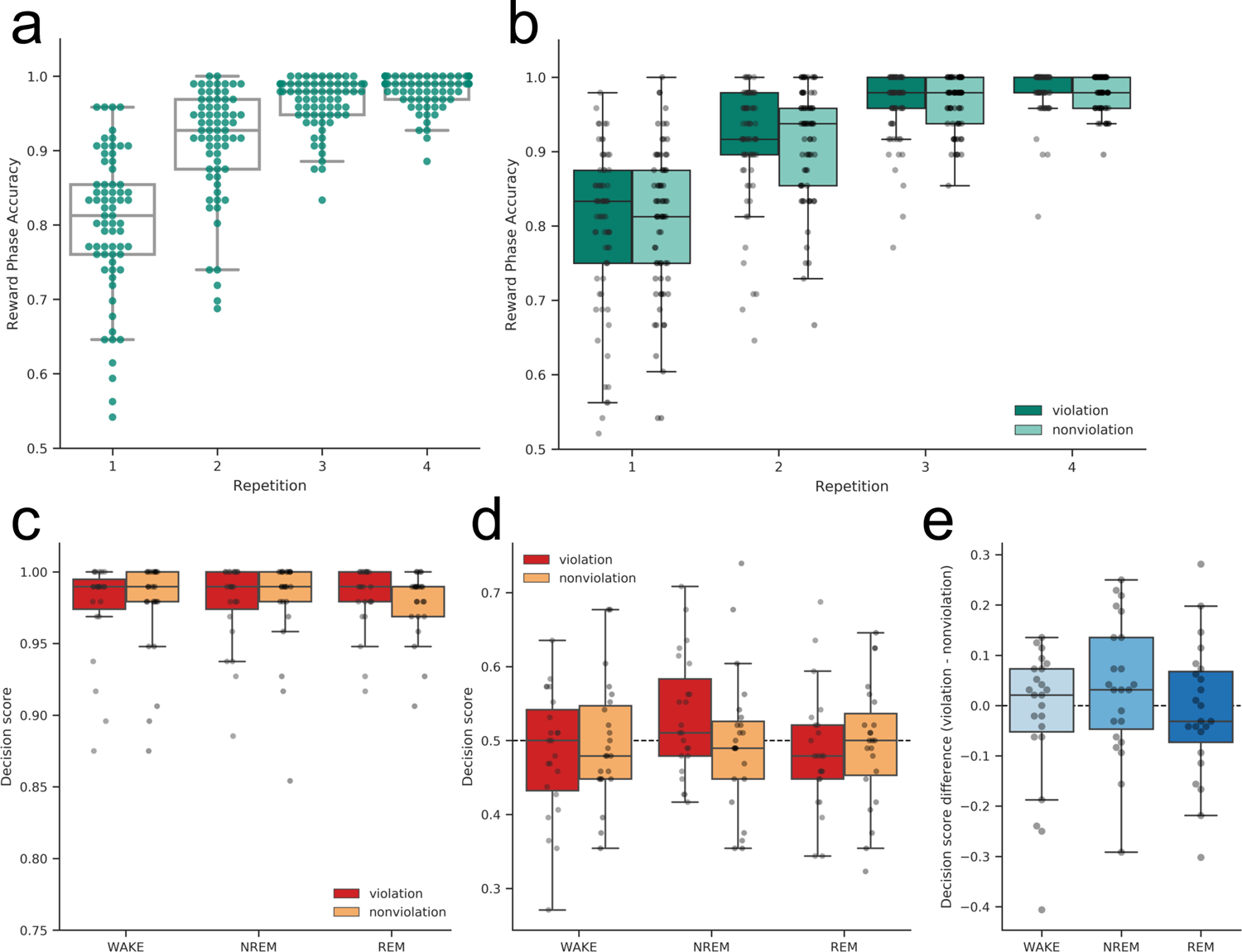
Reward learning task behavioral performance measures. During the reward prediction phase, participants saw an A scene and predicted if that scene was associated with a reward or neutral outcome. (a) Reward prediction accuracy increased across four learning repetitions (*p*< 0.001), and (b) there was no interaction between pair type (violation, nonviolation) and learning repetition. During the decision phase, participants were shown one rewarded and one neutral scene, and chose the one they thought was most likely to lead to winning a reward. (c) The decision score for explicitly-learned A scenes was near 1 for most participants, reflecting the strong learning of A scene reward associations. (d) The decision score on B trials was no different than chance in any group or condition, and (e) no group showed a difference in decision bias between pair types (violation-nonviolation) significantly different from zero.

## Supplementary Table

**Table S1.** Sleep variables correlated with the neural differentiation score.

|  | CA2/3/DG |  |  | CA1 |  |  |
| --- | --- | --- | --- | --- | --- | --- |
|  | left | right | bilateral | left | right | bilateral |
| TST (min) | −0.087 | −0.150 | −0.138 | −0.028 | 0.037 | 0.019 |
| N1 (min) | −0.086 | −0.029 | −0.092 | −0.082 | −0.104 | −0.140 |
| N2 (min) | −0.055 | 0.049 | 0.001 | 0.104 | 0.189 | 0.197 |
| N3 (min) | 0.048 | −0.170 | −0.062 | −0.267 | −0.145 | −0.242 |
| REM (min) | −0.120 | 0.099 | 0.011 | 0.361 | −0.018 | 0.197 |
| N2 spindle density | −0.002 | −0.119 | −0.084 | −0.013 | 0.120 | 0.112 |
| N3 spindle density | −0.011 | 0.170 | 0.116 | 0.362* | 0.158 | 0.336* |

Note: Values reported are Pearson's *r*. Spindle densities are from electrode CPz. All correlations include *n* = 44, except REM minutes is *n* = 22, and N3 spindle density is *n* = 43 (one participant had 0 minutes of N3 sleep). \**p* < 0.05, not corrected for multiple comparisons

## Notes

### Competing Interest Statement

The authors have declared no competing interest.

